# Hi-Compass resolves cell-type chromatin interactions by single-cell and spatial ATAC-seq data across biological scales

**DOI:** 10.1101/2025.05.14.654019

**Authors:** Yuan-Chen Sun, Wen-Jie Jiang, Kang-Wen Cai, Na-Na Wei, Fu-Ting Lai, Rui-Xiang Gao, Ze-Yu Kuang, Jia-Lu Zhou, An Liu, Han-Wen Zhu, Ming Xu, Hua-Jun Wu

**Author notes:** Correspondence (M.X) (H.J.W.). These authors contributed equally (Yuan-Chen Sun, Wen-Jie Jiang, Kang-Wen Cai).

## Abstract

Computational prediction of three-dimensional (3D) genome organization provides an alternative approach to overcome the cost and technical limitations of Hi-C experiments. However, current Hi-C prediction models are constrained by their narrow applicability to studying the impact of genetic variation on genome folding in specific cell lines, significantly restricting their biological utility. We present Hi-Compass, a generalizable deep learning model that accurately predicts chromatin organization across diverse biological contexts, from bulk to single-cell samples. Hi-Compass outperforms existing methods in prediction accuracy and is generalizable to unseen cell types through chromatin accessibility data, enabling broad applications in single cell omics. Hi-Compass successfully resolves cell-type-specific 3D genome architectures in complex biological scenarios, including immune cell states, organ heterogeneity, and tissue spatial organization. Furthermore, Hi-Compass enables integrative analysis of single-cell multiome data, linking chromatin interaction dynamics to gene expression changes across cell clusters, and mapping disease variants to pathogenic genes. Hi-Compass also extends to spatial multi-omics data, generating spatially resolved Hi-C maps that reveal domain-specific chromatin interactions linked to spatial gene expression patterns.

## Introduction

The three-dimensional (3D) organization of the genome is a critical regulator of gene expression, cellular function, and tissue development^1–6^. High-throughput techniques, particularly chromosome conformation capture (3C)-based methods such as Hi-C, have revolutionized our understanding of chromatin architecture by mapping interactions at genome-wide scales^7, 8^. These studies have revealed hierarchical structures, including A/B compartments, topologically associating domains (TADs), and chromatin loops^9–17^, which collectively orchestrate cell-type-specific gene expression through enhancer-promoter interactions. Such insights are essential for deciphering the regulatory mechanisms underlying development and disease^6, 18–21^.

Multiple factors contribute to the formation of 3D genome architecture. DNA sequences serve as the fundamental blueprint, with specific motifs directing nucleosome positioning, protein complex binding, and the establishment of regulatory domains^18, 22, 23^. Chromatin accessibility, which reflects the activity of regulatory elements such as enhancers and promoters, further modulates chromosomal interactions^24–26^. Architectural proteins, including CTCF, play a pivotal role by facilitating chromatin looping and boundary formation^27–30^. The dynamic interplay of these elements creates cell-type-specific 3D organization patterns which are essential for maintaining cellular identity and function^31, 32^.

Despite its transformative impact, Hi-C technology faces practical challenges. Generating high-resolution Hi-C data requires substantial resources, specialized expertise, and extensive sequencing depth^33, 34^. Therefore, computational approaches that predict 3D genome structure have emerged as a valuable and feasible solution. For instance, DeepC^35^ employs transfer learning to predict chromatin interactions from DNA sequences leveraging epigenomic pre-training. Akita uses a convolutional neural network to predict interactions at 2 Kb resolution from 1 Mb DNA sequences, accounting for distance-dependent decay. C.Origami^33^ represents a multimodal architecture integrating DNA sequences, bulk CTCF ChIP-Seq and ATAC-seq signals through CNN layers and self-attention blocks. In contrast, Epiphany^36^ and ChromaFold^37^ predicts contact maps solely from epigenomic information, bypassing DNA sequences, through a much lightweight model.

The rapid expansion of single-cell and spatial epigenomic profiling technologies presents a promising opportunity to extend 3D genome prediction models to these emerging data types. However, existing computational methods face critical limitations when applied to single-cell and spatial datasets: (1) Requirement for jointly profiled ATAC-seq, CTCF ChIP-seq and other epigenomics data from the same biological sample; (2) Limited capacity to generate cell-type-specific predictions for unseen cell types; (3) Failure to account for data sparsity inherent in single-cell and spatial omics; (4) Lack of fine-grained structural prediction supporting downstream analytical applications.

To address these limitations, we introduce Hi-Compass, a deep learning model that significantly advances the prediction of near-optimal, cell-type-specific Hi-C maps. Hi-Compass uniquely leverages (sc)ATAC-seq data as its sole cell type-aware input, combined with DNA sequence and a generalized CTCF binding profile, to infer structural details of 3D genome organization. The model dynamically adjusts to varying sequencing depths through a depth-aware module, enabling robust performance across diverse experimental settings, including bulk, single cell and spatial assays. Hi-Compass uses a multi-cell-type training strategy, enabling zero-shot generalization to unseen cell types. Benchmarking analyses demonstrate that Hi-Compass outperforms existing methods in prediction accuracy for bulk samples. Furthermore, extensive evaluations in single-cell and spatial contexts reveal its ability to precisely resolve cell-type-specific 3D genome structures in complex biological scenarios, including immune cell states, organ heterogeneity and tissue spatial organization. Moreover, Hi-Compass can identify long-range regulatory loops linking disease-associated non-coding variants to their target genes, providing mechanistic insights of pathogenic gene dysregulation. By bridging accessible chromatin profile with 3D genome architecture, Hi-Compass expands the range of biological questions that can be addressed through Hi-C prediction models.

## Results

### Hi-Compass enables Hi-C prediction from multi-depth ATAC-seq data

To address the challenge of predicting three-dimensional genome structure from single-cell chromatin accessibility data, we developed Hi-Compass (**Fig. 1a**), a deep learning model integrates four key inputs: ATAC-seq signal, ATAC-seq sequencing depth, DNA sequence and a generalized CTCF binding profile. This multi-modal input design allows the model to understand both the general principles of genome folding from static genomic features and cell-type-specific genome folding from chromatin accessibility patterns. Hi-Compass is designed to adapt to ATAC-seq data of varying sequencing depths, enabling the generation of high-quality, cell-type-specific Hi-C predictions without requiring additional experimental data.

**Fig. 1.**
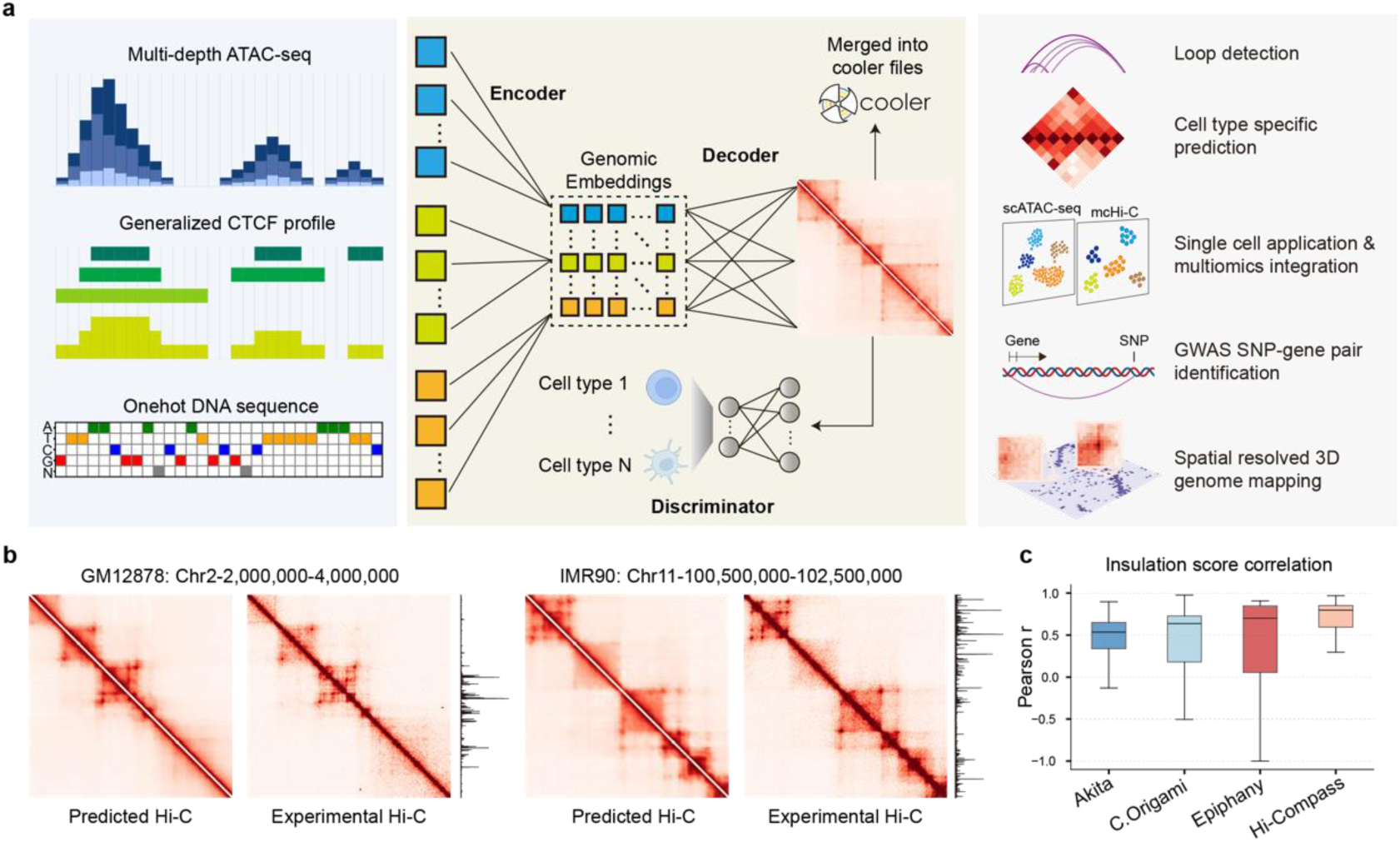
Hi-Compass predicts 3D genome structure in a sequencing depth-aware manner. a, Overview of the Hi-Compass methodology framework. The model takes three types of sequence information as input to predict corresponding Hi-C structures, integrates them into whole-chromosome.cool files, and enables different downstream analysis scenarios for various applications. **b,** Comparison between Hi-Compass-predicted Hi-C and experimental ground truth Hi-C for GM12878 and IMR90 in representative regions of the validation set (Chr2) and test set (Chr11). The plot on the right side of the heatmap depicts the chromatin accessibilities of the corresponding regions. **c,** Performance comparison of Hi-Compass with three previously published Hi-C prediction methods: Akita, C.Origami, and Epiphany. Box plots show the correlation of insulation scores between predictions and experimental Hi-C data. Chromosomes were divided into segments with a step size of 2 Mb to calculate similarity. The insulation scores were uniformly calculated for all methods using a window size of 20 bins, excluding the first two diagonals

Since only matched bulk ATAC-seq and Hi-C data are currently available, we developed a computational strategy to simulate multi-depth ATAC-seq profiles spanning a continuum from single-cell to multi-cell to bulk-level resolution. This approach allows the model to effectively adapt to scATAC-seq of varying depths. To overcome the limited availability of single-cell CTCF ChIP-seq data, we used a generalized CTCF binding profile derived from pan-tissue samples. Specifically, we integrated bulk CTCF ChIP-seq data from 165 human samples (**Supplementary Table S1**) and constructed a comprehensive CTCF binding map through systematic peak integration. This pan-tissue reference map preserves fundamental CTCF binding principles, serving as a framework to guide ATAC-seq signals in identifying cell-type-specific CTCF binding patterns. For DNA sequence, we employed a five-channel one-hot encoding representing A, T, C, G, and N.

The core architecture of Hi-Compass combines CNN and Transformer modules (**Supplementary Fig. 1**). At the input layer, three encoders composed of convolutional layers extract features from DNA sequences, ATAC-seq signals, and generalized CTCF binding signals. These features are integrated to provide a comprehensive representation of genomic information. The ATAC encoder includes a scaling module that adjusts for total sequencing depth, enhancing the model’s adaptability to varying depths through learnable parameters. The decoder employs a Transformer architecture to process the extracted genomic features and generates the final two-dimensional Hi-C matrix. To improve cell-type-specific prediction accuracy, Hi-Compass incorporates a discriminator network that evaluates the cell-type specificity of the output. The loss function combines mean squared error (MSE) and information-weighted structural similarity^38, 39^ (SSIM) to supervise numerical accuracy and structural features, along with the discriminator’s cross-entropy (CE) to optimize cell-type specificity.

Hi-Compass predicts Hi-C fragments of 2 Mb in size at 10-kb resolution. By sliding the prediction window along chromosomes and adopting an optimized stitching strategy, we built a complete pipeline (**Fig. 1a**) capable of generating genome-wide Hi-C map predictions. These predictions are integrated into standard cool format files for subsequent analyses, such as insulation score calculation and loop identification, by using standard tools.

We curated 14 high-quality matched Hi-C and ATAC-seq datasets (**Supplementary Table S2**) covering multiple tissue types, of which 6 samples were used for model training and the rest for testing (**Supplementary Table S3**), providing a diverse set for the model to learn and test. In determining the optimal threshold of ATAC-seq sequencing depth, we systematically explored its impact on signal reservation. Depth reduction preserves most genomic features from bulk sequencing profiles (**Extended Fig. 1a**), with high reproducibility observed between different sampling replicates at equivalent depths (**Extended Fig. 1b,c**) until reaching critical depth threshold. Based on these assessments, we implemented a minimum ATAC-seq depth threshold of 2e5 reads (∼ 0.01X genome coverage) during model training to ensure robust and specific predictions. For rigorous evaluation, we designated chromosome 2 as the validation set and chromosome 11 as the test set. Hi-Compass predictions reproduced the topological structural details of experimental Hi-C with high fidelity in both validation and test (Fig. 1b).

### Systematic evaluation of Hi-Compass performance

We systematically compared Hi-Compass with state-of-the-art Hi-C prediction methods, including Akita^40^, C.Origami^33^ and Epiphany^36^. To ensure a fair evaluation, we used identical test datasets and assessment metrics for all methods. Insulation score^41^ correlation was applied as the primary evaluation metric due to its ability to effectively capture chromatin structural features and its robustness to batch effects^33^.

Hi-Compass significantly outperformed other methods in insulation score correlation with experimentally measured Hi-C data (**Fig. 1c**). In genome-wide evaluations based on 2 Mb windows, Hi-Compass achieved a median correlation coefficient above 0.7, while other methods fell significantly below this threshold and exhibited less stability. To visually demonstrate the performance of Hi-Compass, we selected multiple representative genomic regions and compared Hi-Compass predictions side-by-side with those from existing methods (**Extended Fig. 2**). These regions cover diverse chromatin structural features, including TAD boundaries and local interaction patterns. Visual comparisons clearly revealed that Hi-Compass-generated Hi-C matrices closely resembled experimentally measured Hi-C data, particularly in capturing fine-scale structural details. In contrast, while other methods captured major structural features, they often exhibited loss of microstructure details, artificial segmentations and blurred profiles.

### De novo inference of cell-type-specific Hi-C maps with visible chromatin loop structures

To demonstrate the generalizability of Hi-Compass, we evaluated its prediction performance on multiple unseen samples. Using bulk ATAC-seq data from diverse samples, we predicted their genome-wide Hi-C maps and calculated the Pearson correlation of insulation scores between predictions and experimental data. Hi-Compass predictions exhibited strong cell-type specificity, with high correlations along the diagonal of the heatmap (**Fig. 2a and Extended Fig. 3a**). We then conducted distance-stratified correlation analysis, calculating the correlation between predictions and experimental data at varying genomic distances (**Extended Fig. 4**). Across all tested samples, Hi-Compass exhibited stable prediction accuracy.

**Fig. 2.**
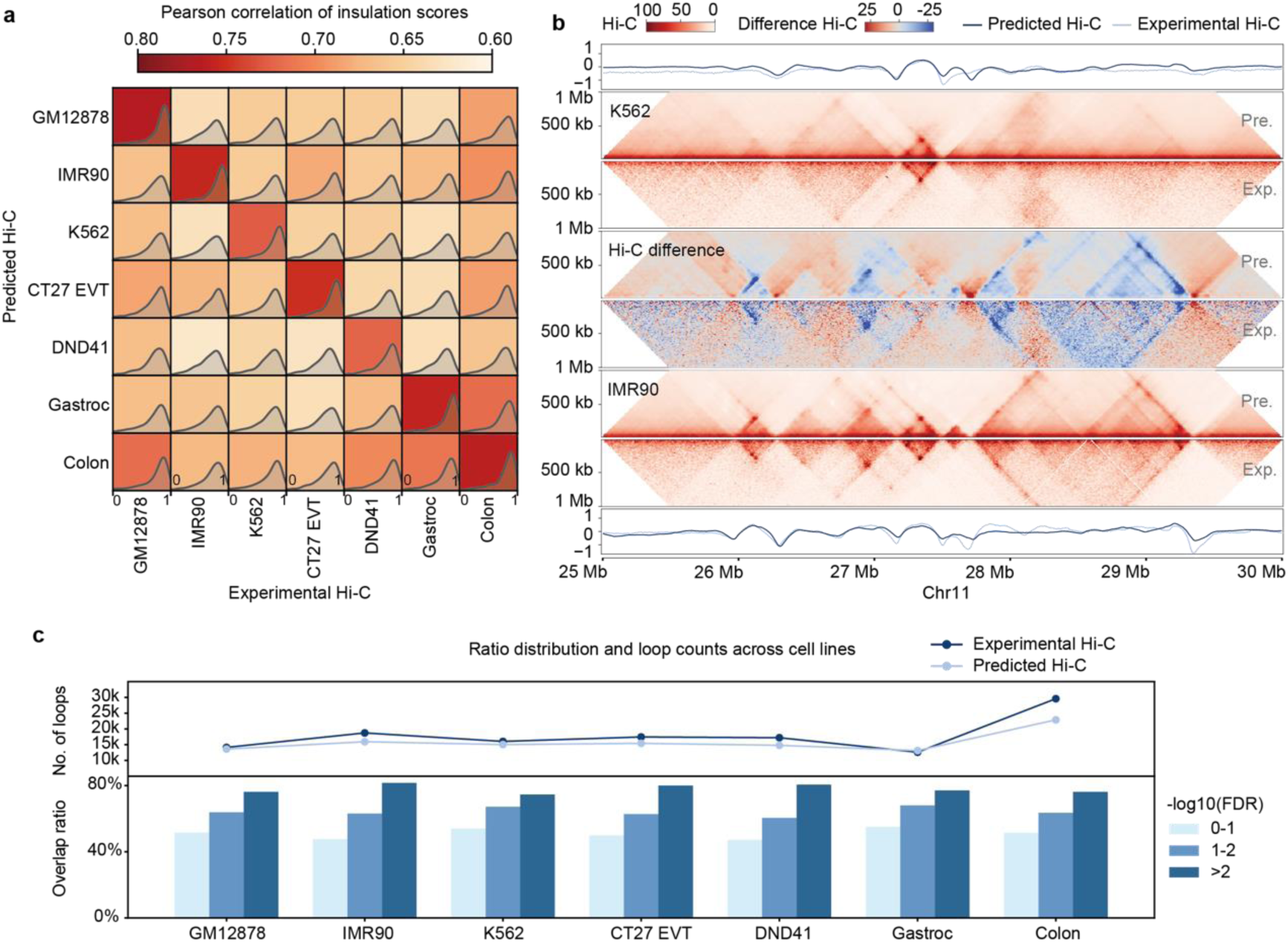
De novo inference of cell-type-specific Hi-C maps with visible chromatin loop structures. a, Validation of chromatin domain prediction accuracy through insulation score correlation analysis. Heatmap displays Pearson correlation coefficients of insulation scores between Hi-Compass predictions and experimental Hi-C data across multiple samples. The first three are from the training set, and the last four are from the test set. Insulation scores were calculated using a window size of 20 bins, excluding the first two diagonals of the Hi-C matrix. Gastroc: Gastrocnemius, Colon: Transverse colon. **b,** Comparative analysis of predicted and experimental Hi-C in GM12878 (top) and IMR90 (bottom) cells. The heatmap in between shows the difference between these two cell lines in both prediction and experiment. Within each plot, the upper half shows the predicted results and the lower half shows the experimental results. **c,** Quantitative assessment of chromatin loop prediction performance. Top: Comparison of total predicted versus experimentally observed loops across multiple samples. Bottom: Precision analysis showing the fraction of gold-standard loops captured by predictions, stratified by FDR thresholds in experimental data.

Visual comparisons of Hi-C maps revealed that Hi-Compass predictions not only achieved high overall accuracy but also precisely preserved cell-type-specific structural features (**Fig. 2b**). The predicted maps closely matched the corresponding experimental data, accurately capturing key features such as compartmental structures, TAD boundaries, and chromatin loops. Differential contact analysis between distinct cell lines further highlighted the cell-type-specific TADs and loops captured by Hi-Compass closely align with experimental observations.

The high-quality predictions of Hi-Compass enable robust downstream analyses. Loop detection on both Hi-Compass-predicted and experimental Hi-C data revealed strong concordance, particularly for highly significant loops (**Fig. 2c and Extended Fig. 3b**). At a stringent FDR cutoff (FDR<0.01), Hi-Compass recovered approximately 80% of experimentally identified loops. Additionally, we analyzed the overlap between Hi-Compass-predicted loops and experimentally identified ones^18, 42^, revealing significant consistency (**Extended Fig. 3c**). For example, in DND41, 10,534 of 14,793 predicted loops overlapped with experimental ones (71% consistency), while in Gastrocnemius, 8,345 of 13,130 predicted loops overlapped with experimental ones (64% consistency). The average loop prediction consistency was 63.9%, approaching the 67.1% reported between high-quality experimental replicates of GM12878^18^. This performance demonstrates that Hi-Compass can reliably resemble chromatin interactions across diverse cell types, close to the experimental replicates. Considering the inherent challenges of loop detection itself, these results underscore the practical utility of Hi-Compass predictions for guiding experimental validation and functional studies.

### Hi-Compass overcomes single-cell data sparsity for robust Hi-C prediction

Building on Hi-Compass’s high performance with bulk data, we extended its application to single-cell settings. Leveraging insights from our ATAC-seq depth analysis (**Extended Fig. 2**), we implemented a meta-cell strategy to address the inherent sparsity of single-cell data (**Extended Fig. 5a**). Specifically, we integrated ATAC-seq signals from adjacent single cells into meta-cells, ensuring that the input data met the depth threshold required for accurate, cell-type-specific predictions.

We applied Hi-Compass to a scATAC-seq dataset comprising seven cell lines (**Supplementary TableS4**), four of which (IMR90, GM12878, K562, and HCT116) had corresponding bulk Hi-C data available (**Fig. 3a**). Meta-cell Hi-C (mcHi-C) was predicted from meta-cell ATAC-seq, with each meta-cell containing an average of 10 cells (range: 5-15). We extracted the first 50 diagonals from each predicted Hi-C matrix as feature vectors, performed PCA dimensionality reduction, and visualized the results using UMAP. The mcHi-C predictions from the same cell line formed distinct clusters (**Fig. 3b**), confirming the high consistency and cell type specificity of Hi-Compass predictions. Additionally, the Hi-C predictions for each meta-cell showed clear chromatin structures and cell-type-specific patterns across the genome, as illustrated in a representative genomic region (**Fig. 3c**).

**Fig. 3.**
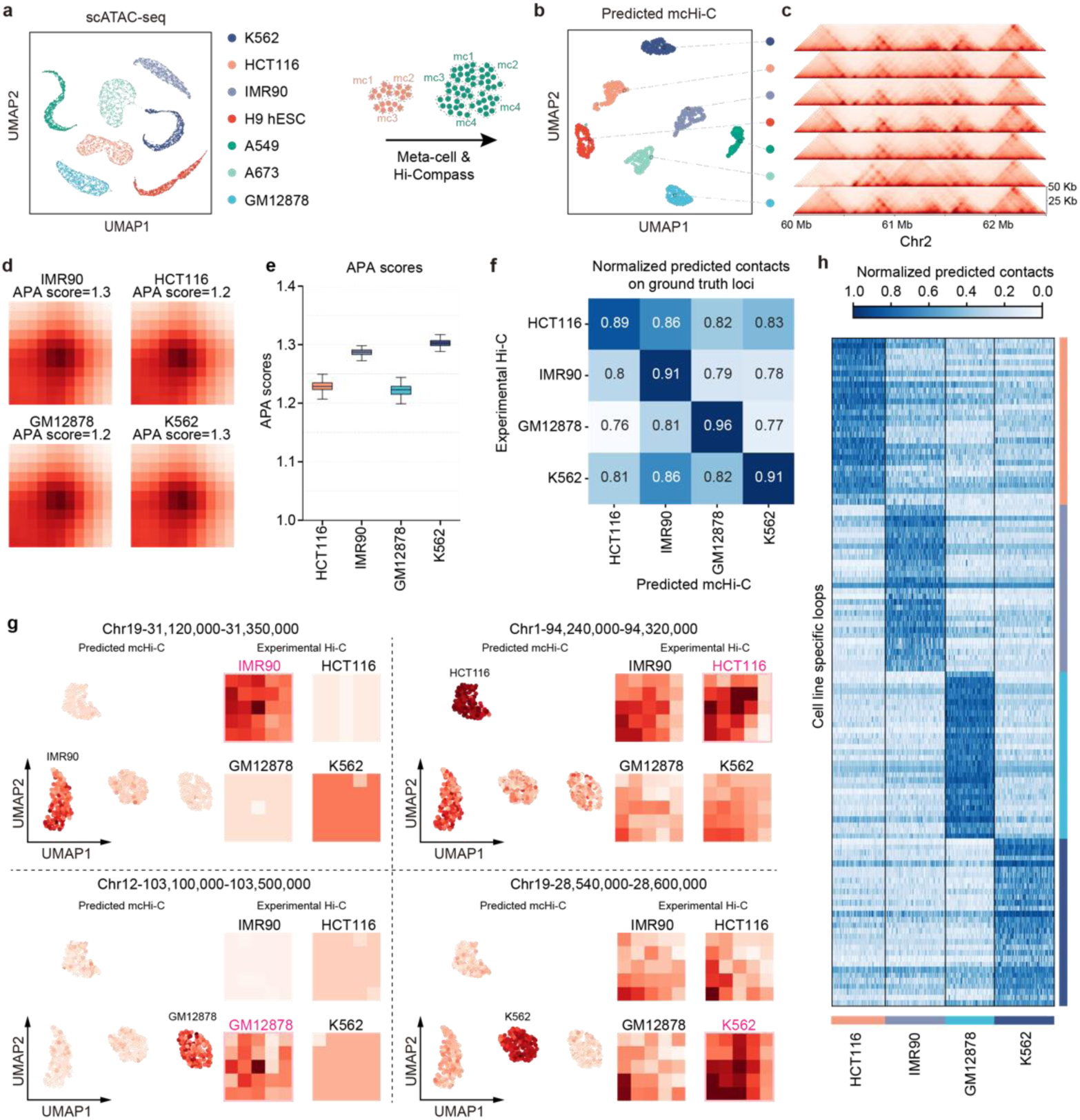
Hi-Compass predicts cell-type-specific 3D chromatin interactions using scATAC-seq. a. UMAP projection of scATAC-seq profiles of seven cell lines. **b.** UMAP visualization of predicted mcHi-C. **c.** Example predicted mcHi-C on a genomic region from Chr2. Lateral Hi-C plot showing one meta cell profile per cell type. **d.** APA plots show the aggregate signals of predicted mcHi-C from experimentally detected loops in four cell lines (IMR90, HCT116, GM12878, K562). The plots are generated by summing submatrices of predicted contacts of mcHi-C surrounding loop peaks detected in experimental Hi-C data. **e.** Boxplot showing the distribution of APA scores per meta cell across cell lines. **f.** Cross-validation of cell-type-specificity depicted by normalized predicted contacts on experimentally detected loop peaks across cell lines. **g.** Representative cell line-specific loops visualized by feature plots of predicted mcHi-C data and loop signal plots of experimental Hi-C data in four cell lines. Each panel shows one cell line-specific chromatin loop as example. Feature plot (left) displays the normalized predicted contacts at the loop peak in each of all meta cells. Loop signal plot (right) displays the experimental Hi-C signals at the same loop peak. **h.** Heatmap displays normalized predicted contacts at cell line-specific loop peaks. Each row represents an individual loop peak, while columns represent predicted contacts at loop peaks across different cell lines.

To quantitatively assess Hi-Compass’s performance, we employed aggregate peak analysis^18^ (APA). First, we called chromatin loops in the four matched experimental Hi-C datasets. These loop coordinates were then mapped to Hi-Compass predictions, and local contact matrices were extracted around each loop peak. APA profiles were generated by superimposing these matrices, revealing significant signal enrichment at actual loop peaks, with contact frequencies at the center point markedly higher than in surrounding regions (**Fig. 3d**). These results closely matched the APA profiles derived from experimental Hi-C data (**Extended Fig. 5b**). Importantly, each independent mcHi-C prediction achieved strong APA scores (APA score > 1) (**Fig. 3e**), with high consistency across meta-cell predictions from the same cell line.

Next, we sought to investigate whether Hi-Compass could accurately predict cell-type-specific chromatin loops. APA analysis was performed on cell-type-specific loops in the four cell lines, revealing that predicted profiles exhibited the strongest signals at their corresponding true loop peaks (**Fig. 3f and Extended Fig. 6a**). Predicted loop peak signals were highly consistent within the same cell line and aligned well with experimental data, as illustrated by representative loops (**Fig. 3g and Extended Fig. 6b**). Further analysis revealed that most of experimentally identified cell-type-specific loops were precisely captured by mcHi-C predictions with statistical significance (effect size > 0.6, p < 0.05) (**Fig. 3h and Extended Fig. 6c**). These results collectively demonstrate Hi-Compass’s reliability and stability in predicting cell-type-specific chromatin structures from sparce scATAC-seq data, highlighting its potential for broader applications in single-cell 3D genomic research.

### Hi-Compass reconstructs cell type chromatin dynamics in complex tissues

To explore the application potential of Hi-Compass in complex tissues, we analyzed a peripheral blood mononuclear cells (PBMC) scATAC-seq dataset^43^. Cell type annotations were assigned using label transfer from an annotated PBMC scRNA-seq dataset. After filtering out cell types with insufficient cell numbers, we retained 10 major immune cell subtypes for subsequent analysis (**Fig. 4a**). A similar meta-cell strategy followed by Hi-Compass were applied. The predicted mcHi-C effectively preserved the hierarchical relationships of immune cell types, as visualized by UMAP. Meta-cells within the same cell type exhibited strong correlations while maintaining distinct separation from other cell types (**Fig. 4b**). These results were not biased by variations in ATAC-seq sequencing depth across meta-cells (**Supplementary Fig. 2a**).

**Fig. 4.**
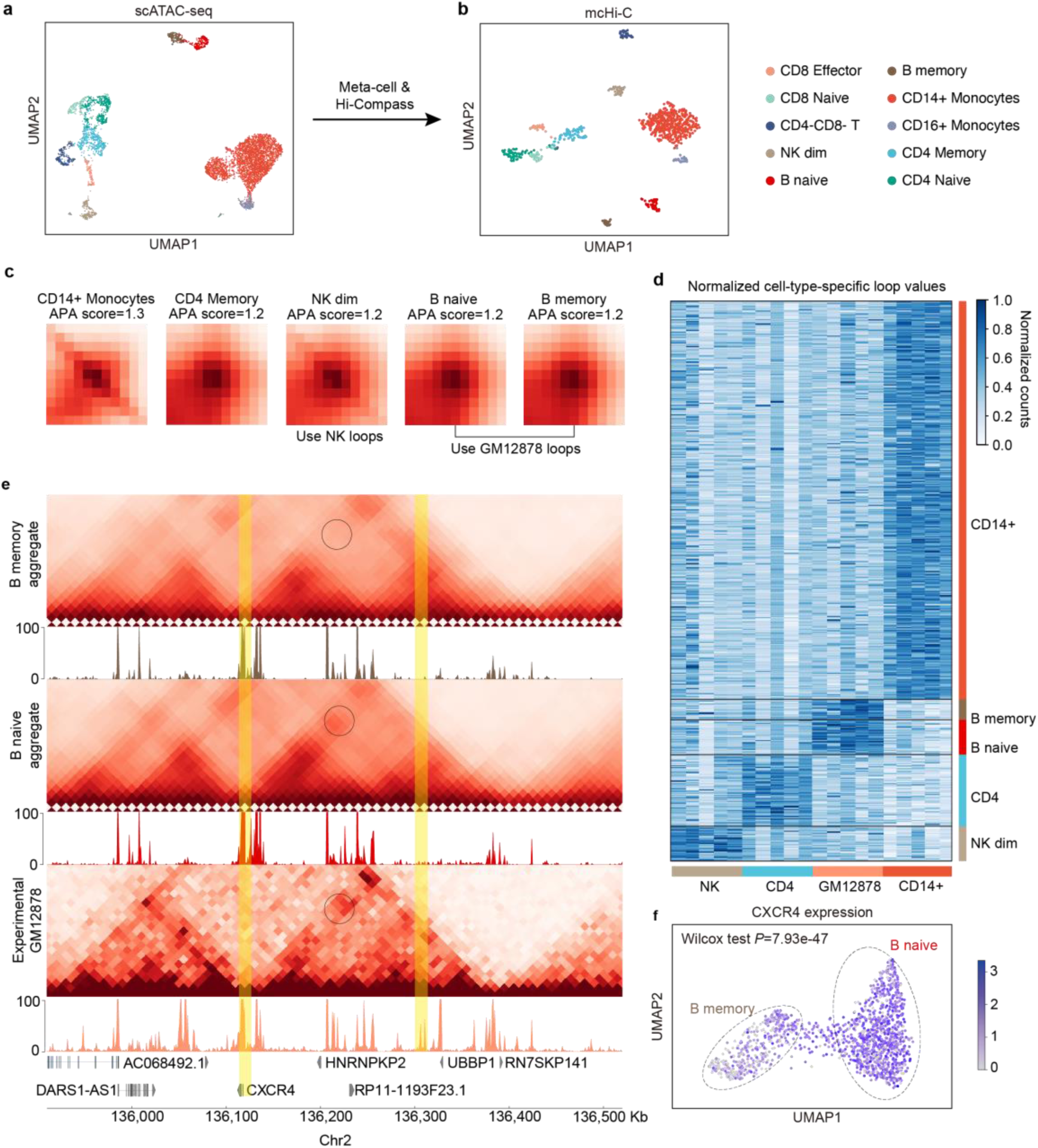
Hi-Compass generates accurate cell-type-specific prediction in complex tissues. **a**, UMAP projection of PBMC scATAC-seq profiles, with different colors representing immune cell subtypes. **b**, UMAP visualization of predicted mcHi-C. **c**, APA plots show the aggregate signals of predicted mcHi-C from experimentally detected loops in four representative immune cell types (CD14+ Monocytes, CD4 Memory, NK dim, and B progenitor). The plots are generated by summing submatrices of predicted contacts of mcHi-C surrounding loop peaks detected in experimental Hi-C data. **d**, Heatmap displays normalized predicted contacts at cell-type-specific loop peaks. Each column represents an individual loop peak, while rows represent predicted contacts at loop peaks across different cell types. **e**, Predicted Hi-C profiles and corresponding ATAC-seq signal tracks for B cell subtypes and GM12878 in the CXCR4 region of Chr2. The loop anchored at CXCR4 TSS is marked in black. Loop anchors are highlighted in yellow. **f**, Feature plot showing CXCR4 expression levels within two B cell subtypes.

To evaluate prediction accuracy, we collected publicly available bulk Hi-C data for four key immune cell types: CD4 Memory T cells, CD14+ monocytes, NK cells (corresponding to NK dim), and GM12878 (corresponding to B naive and B memory cells). These verification datasets encompassed major immune cell lineages and diverse functional characteristics, providing an ideal reference for assessing Hi-Compass’s performance in complex immune microenvironments. APA analysis revealed significant enrichment of predicted mcHi-C signals at true loop peaks across immune cell subtypes (**Fig. 4c and Supplementary Fig. 2b**), with APA scores consistently exceeding 1 for all meta cells (**Extended Fig. 7a,b**). Further analyses unveiled that Hi-Compass accurately recovered cell-type-specific loop signals (**Fig. 4d and Extended Fig. 7c,d**).

To elucidate the capability of Hi-Compass in discovering 3D genome dynamics in closely related cell states, we delved into the predicted mcHi-C profiles of two B cell states: B memory and B naïve. GM12878 experimental data served as a reference. We focused on dynamic chromatin reorganization around key factors in B cell differentiation. Hi-Compass predictions identified a cell state-specific chromatin loop connecting the CXCR4 promoter with a distal regulatory element in naïve B cells that was absent in memory B cells (**Fig. 4e**). Notably, while chromatin accessibility at the regulatory region showed subtle differences between these cell states, the presence of this Hi-C interaction in naïve B cells correlated strongly with significantly higher CXCR4 expression levels (**Fig. 4f and Supplementary Fig. 2c**), validating the functional relevance of the predicted conformation change. These results demonstrate Hi-Compass’s sensitivity in detecting fine-scale but biologically significant 3D genome reconfigurations that underlie gene expression differences in closely related cell states.

As a critical chemokine receptor involved in B cell development, homing, and tissue distribution, CXCR4 exhibited cell-state-specific chromatin conformation changes that likely reflect its fine-tuned transcriptional regulation^44–46^. This multi-omics integrated analysis not only showcases Hi-Compass’s utility in addressing biological questions but also demonstrates its capability to reliably predict 3D genomic structures from scATAC-seq data in novel cell states.

### Hi-Compass integrates single-cell multiome data to reveal functional associations

In this section, we applied Hi-Compass to a single-cell multiome dataset that sequenced a 105-day female human embryonic heart tissue. The data contains jointly profiled gene expression and chromatin accessibility in the same single cells. We processed the data using standard procedures, and annotated cell types based on gene expression profiles. Unsupervised clustering of scATAC-seq data revealed high consistency with transcriptome-annotated cell classifications (**Fig. 5a**), confirming the internal coherence of the dataset. Following the previous workflow, we generated mcHi-C predictions from scATAC-seq data. UMAP dimensionality reduction demonstrated that predicted mcHi-C maintained distribution patterns similar to scATAC-seq data (**Fig. 5b**), indicating that Hi-Compass successfully captured chromatin conformation characteristics across different cell types. We then identified differential loops for each cell type and integrated gene expression data from the same cells to validate the biological relevance and reliability of Hi-Compass predictions.

**Fig. 5.**
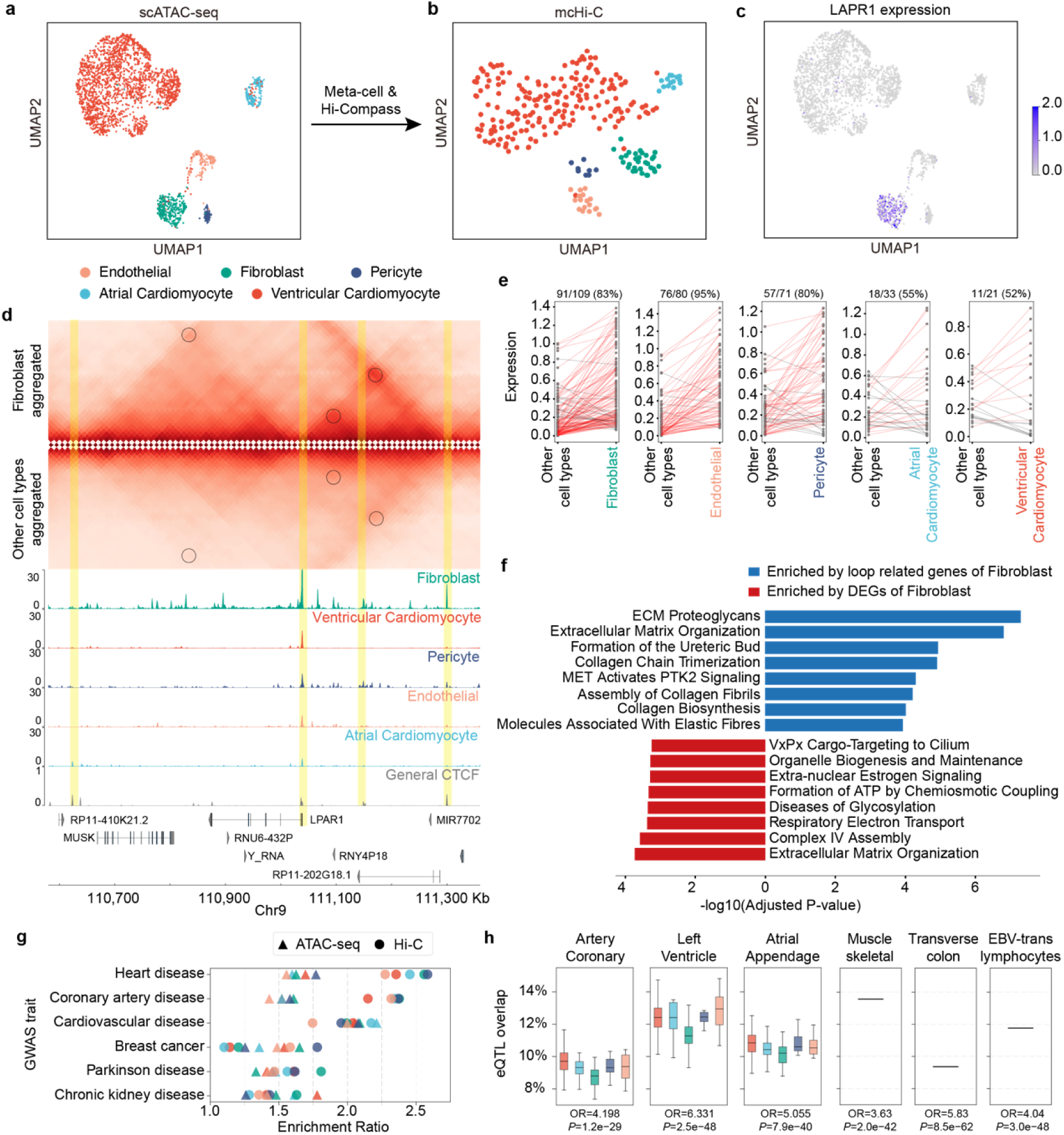
Hi-Compass integrates single-cell multiome data to reveal functional associations. **a**, UMAP projection of scATAC-seq profile from human embryonic heart tissue, with different colors representing distinct cell types. **b**, UMAP visualization of predicted mcHi-C, demonstrating the prediction reserved the original cell type distribution pattern. **c**, Feature plot showing LPAR1 is specifically expressed in fibroblasts. **d**, Predicted Hi-C profiles and corresponding ATAC-seq signal tracks for fibroblasts versus other cell types in the LPAR1 genomic region. The loops anchored at LPAR1 TSS are marked in black. Loop anchors are highlighted in yellow. **e**, Genes associated with cell-type-specific loops are frequently upregulated in corresponding cell types (marked by red lines). Genes whose promoters located in a cell-type-specific loop anchor are included in the analysis. **f**, Reactome pathway enrichment analysis of upregulated DEGs (red) and differential loop-related genes (blue) in fibroblasts. **g**, Enrichment analysis of GWAS variants across disease categories. Dot plot showing enrichment scores for cardiac versus non-cardiac traits, comparing SNP annotations based on: Hi-Compass-predicted loop anchors (triangular) versus ATAC-seq peaks (circular). Cardiac disease categories show significantly stronger enrichment for loop-annotated variants. **h**, Validation of SNP-gene regulatory links against GTEx eQTL data. Box plots showing overlap percentages between loop-annotated SNP-gene links and eQTL data in different tissues. The left three panels display results using Hi-Compass-predicted Hi-C data, while the right three show results derived from experimental bulk Hi-C data.

Taking fibroblast as an example, a marker gene LPAR1 exhibited significantly higher expression in fibroblasts compared to other cell types (**Fig. 5c**). By comparing Hi-Compass predictions between fibroblasts and other cell types, we identified fibroblast-specific chromatin loops anchored at the LPAR1 TSS (**Fig. 5d**). These loops were supported by cell-type-specific ATAC peak signals, with chromatin accessibility at loop anchors significantly higher in fibroblasts. Similar biologically interpretable loops were predicted across other cell types. For instance, in endothelial, the PTPRM gene region displayed cell-type-specific chromatin interactions (**Extended Fig. 8a**), consistent with its high expression in these cells (**Extended Fig. 8b**). Similarly, atrial cardiomyocytes exhibited cell-type-specific chromatin conformations in the MAPK4 gene region (**Extended Fig. 8c**), corresponding to its cell-type-specific expression pattern (**Extended Fig. 8d**). Systematic analysis of all cell-type-specific loops revealed that genes associated with these loops generally exhibited higher expression in their corresponding cell types (**Fig. 5e**). This observation highlights the coordinated relationship between cell-type-specific chromatin structures and gene expression regulation, further validating the biological relevance of Hi-Compass predictions.

By comparing the overlap of predicted mcHi-C loops across cell types, we identified unique chromatin interaction patterns that may indicate functional associations (**Supplementary Fig. 3a**). The shared loops between fibroblasts and pericytes were the second most abundant, surpassed only by those between two cardiomyocyte types, suggesting potential interconnected regulatory networks between the two cell types in embryonic heart development. Pathway analysis of differentially expressed genes (DEGs) in fibroblasts and fibroblast-specific chromatin loop-associated genes (loop genes) both revealed extracellular matrix (ECM) organization as the top enriched pathway (**Fig. 5f**). Intersectional analysis identified 22 genes common across developmental stages, including critical extracellular matrix components (COL14A1, FBN1), transcriptional regulators (TBX18, GLI2, GLIS3), and cell migration and adhesion molecules (DCC, CDH19) (**Supplementary Fig. 3b**). Notably, loop genes uniquely enriched for fiber-associated pathways, suggesting direct cell identity regulation through 3D genome organization. These results demonstrate Hi-Compass’s ability to discover novel chromatin structures and regulatory relationships in single-cell multiome datasets.

### Hi-Compass links disease variants to pathogenic genes

Beyond identifying cell-type-specific regulatory networks, we explored the potential of Hi-Compass for translating epigenomic information into clinically relevant insights. Using predicted chromatin interactions from embryonic heart tissue, we systematically analyzed SNP-to-gene regulatory relationships by connecting non-coding regulatory elements with gene promoters through predicted loops. Remarkably, enrichment analysis of Hi-Compass-predicted loop anchors on GWAS variants revealed striking specificity for cardiac-related disorders (**Fig. 5g**), with heart disease, coronary artery disease, and cardiovascular disease showing significantly higher enrichment than non-cardiac ones. Crucially, these loop-annotated variants demonstrated substantially greater enrichment than ATAC-seq peak-annotated ones for cardiac traits, highlighting a new perspective of using Hi-Compass-predicted loops to prioritize pathogenic non-coding variants in tissue related diseases.

We next compared Hi-Compass-predicted SNP-to-gene links with tissue-specific expression quantitative trait loci (eQTLs) from the Genotype-Tissue Expression (GTEx) project. In artery coronary, left ventricle, and atrial appendage, Hi-Compass-predicted links showed an average of 9%, 13%, and 11% overlap rates with eQTLs, respectively (Fig. 5h). These numbers are very close to the results obtained by using experimental bulk Hi-C, confirming the high fidelity of our predictions.

Further investigation revealed specific cases of cardiac disease-associated variants physically connected to known heart disease-related genes ^47–51^ through Hi-Compass-predicted loops (**Extended Fig. 9**). We identified regulatory connections linking GWAS variants to established cardiac disease genes, including CDK8, FLRT2, and SSPN. These SNP-to-gene regulatory relationships are supported by both Hi-Compass-predicted chromatin interactions and heart tissue eQTL evidence, suggesting potential role of these non-coding variants in cardiac pathology. These findings demonstrate how Hi-Compass can uncover the regulatory networks underlying disease-associated variants, thereby bridging the gap between genetic association studies and mechanistic understanding of molecular pathology.

### Hi-Compass reconstructs spatially resolved Hi-C maps

Spatial omics technologies have rapidly developed in recent years^52–54^, with spatial transcriptomics making significant progress^55, 56^ and spatial ATAC-seq emerging as a powerful tool^57, 58^. However, the lack of spatial 3D genomic technologies has hindered our understanding of spatially organized chromatin regulatory networks. Here, we explore the potential of Hi-Compass to predict spatially resolved 3D Hi-C maps from spatial ATAC-seq data.

We used a spatial ATAC-RNA co-sequencing dataset from an adult male hippocampus^58^ (**Fig. 6a**) for illustration. Within this dataset, we identified two key regions with clear spatial patterns: the granular layer (ATAC cluster A3, RNA cluster R4) and the choroid plexus (ATAC cluster A5, RNA cluster R6) (**Fig. 6b** and **Fig. 6c**). A major challenge in spatial ATAC-seq is its relatively low sequencing depth, which falls below the input requirement of Hi-Compass. To overcome this limitation, we developed a spatial signal integration strategy similar to the meta-cell approach: merging ATAC-seq data from adjacent areas into “meta spots” to enhance local signal intensity. We then applied Hi-Compass to predict Hi-C for each meta spot, generating spatially resolved chromatin interaction maps for the human hippocampus.

**Fig. 6.**
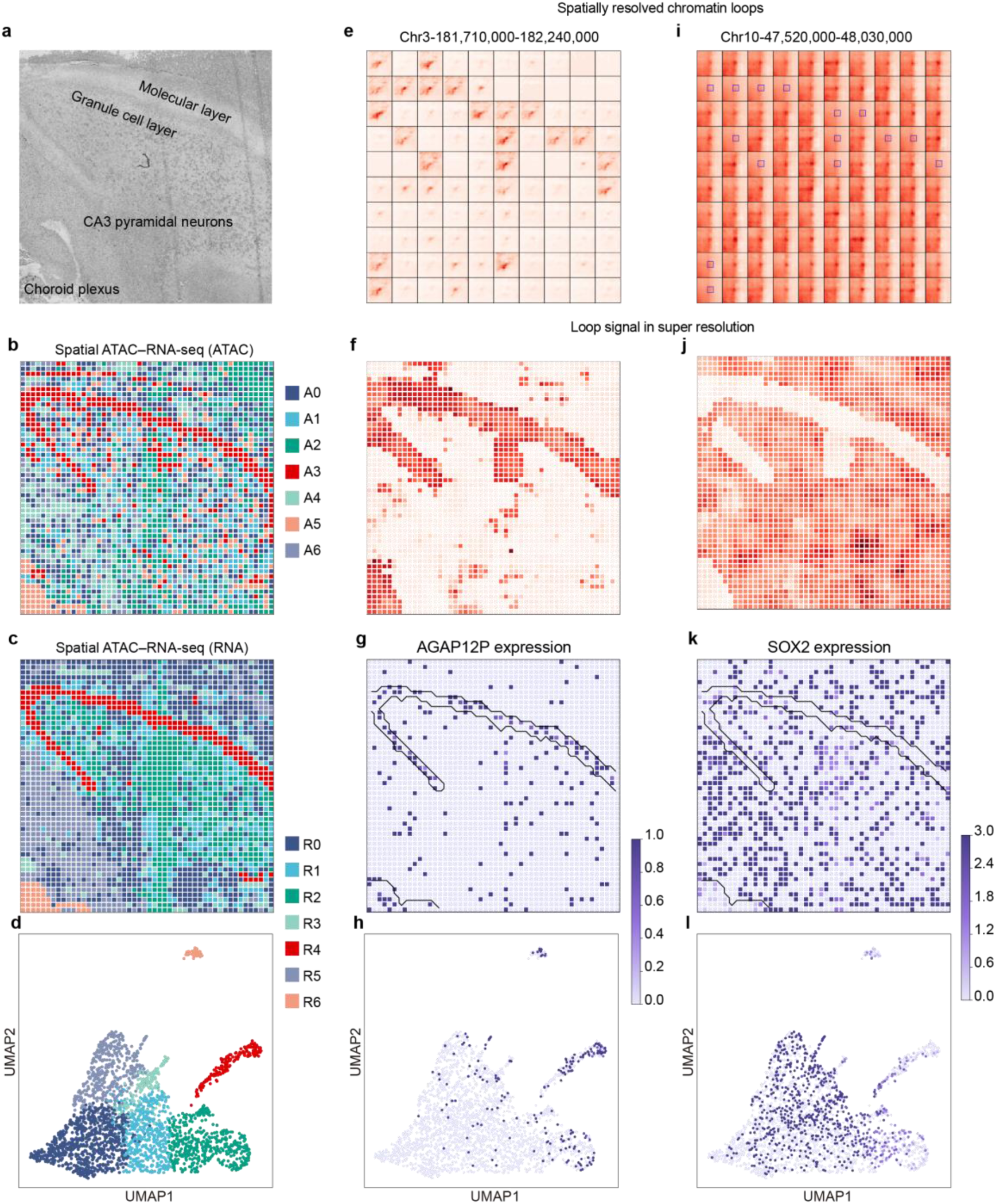
Hi-Compass reconstructs spatially resolved Hi-C maps. **a**, H&E staining image of an adult male hippocampal tissue section, displaying histological characteristics of the granular layer, molecular layer, and choroid plexus. **b, c**, Spatial distribution of ATAC-seq (**b**) and RNA-seq (**c**) clusters, with different colors representing distinct ATAC (A0-A6) and RNA (R0-R6) clusters. Granular layer (ATAC: A3, RNA: R4) and choroid plexus (ATAC: A5, RNA: R6) showing clear region-specific signals and high spatial consistency. **d**, UMAP visualization of spatial RNA clusters. **e,i**, Spatially resolved chromatin loops anchored at AGAP12P (**e**) and SOX2 (**i**) gene TSS regions, obtained from predicted meta-spot Hi-C, demonstrating region-specific chromatin interactions. **f,j**, Super-resolution spatial Hi-C predictions at the AGAP12P (**f**) and SOX2 (**j**) anchored loops. **g,k**, Spatial expression patterns of AGAP12P (**g**) and SOX2 (**k**) genes, with color intensity indicating expression levels. **h,l**, UMAP visualization of AGAP12P (**h**) and SOX2 (**l**) gene expression.

Analysis of the prediction results revealed distinct chromatin conformations in the granular layer and choroid plexus compared to other hippocampal regions (**Extended Fig. 10a,b**). AGAP12P (**Fig. 6e**) and SOX2 (**Fig. 6i**) exhibited region-specific 3D chromatin interactions anchored at their promoters (**Extended Fig. 10c,d**). Their expression levels matched the spatial distribution of predicted chromatin interactions (**Fig. 6g,k,h,l**), suggesting a mechanism that 3D chromatin structure regulates region-specific gene expression in hippocampus.

To enhance spatial resolution and more intuitively demonstrate these differential interactions, we employed a “super-resolution” computational method. By applying the meta spot strategy to each original spot, we generated Hi-C predictions at the same resolution as the original spatial ATAC-RNA data (**Fig. 6f,j**). This approach allowed us to map chromatin interactions while preserving the original spatial resolution. The super-resolution results clearly revealed opposite distribution trends in chromatin interactions anchored at the AGAP12P and SOX2 promoters across different hippocampal regions.

These results demonstrate the feasibility of computationally predicting spatially resolved 3D chromatin structures from spatial epigenomic data, providing novel approach to investigate spatial gene regulation mechanism in complex tissues. Although current spatial ATAC-seq datasets are limited, ongoing advances in spatial epigenomic sequencing technologies will expand the applicability of Hi-Compass to broader tissue types and pathological states. This will offer crucial insights into the relationship between chromatin structure and gene regulation in the spatially resolved manner.

## Discussion

In this paper, we developed Hi-Compass, a generalizable model in predicting 3D chromatin architecture from epigenomic data. Our approach tackles several critical limitations of existing methods by requiring only ATAC-seq signal as experimental input, implementing a meta-cell strategy to overcome single-cell data sparsity, and incorporating a depth-aware module that dynamically adjusts to varying sequencing depths. These designs collectively enable Hi-Compass to generate near-optimal, cell-type-specific Hi-C predictions across diverse biological contexts.

The superior performance of Hi-Compass compared to previous methods is evident in both quantitative metrics and visual inspections of predicted Hi-C matrices. While existing approaches have made important contributions, they typically depend on multiple input modalities and struggle with the inherent data sparsity when generalized to unseen samples and single-cell data. Hi-Compass overcomes these challenges, and effectively captures both local and global chromatin interaction patterns even from limited data inputs. The high correlation between our predictions and experimentally measured Hi-C data, particularly in the identification of topologically associating domains (TADs) and chromatin loops, validates the biological relevance of our predictions.

Besides basic prediction tasks over bulk-level data, Hi-Compass demonstrates remarkable versatility across different scales of biological organizations. At the single-cell level, our meta-cell strategy successfully reconstructs cell-type-specific chromatin interactions from scATAC-seq data, as evidenced by the clear clustering patterns observed in dimensionality reduction analyses and the high concordance of predicted loops with experimentally validated interactions. In complex tissue environments, such as PBMCs, Hi-Compass accurately captures lineage-specific chromatin conformations, enabling the identification of regulatory features that distinguish closely related cell states. Our finding that CXCR4 gene regulation differs between B cell differentiation through differential looping structures exemplifies the biological insights that can be gained through this approach.

The multi-omics integration capabilities of Hi-Compass further extend its utility for biological discovery. By combining predicted Hi-C structures with gene expression data in embryonic heart tissue, we identified cell-type-specific regulatory networks with functional relevance. The discovery that chromatin loops in fibroblasts specifically associate with fiber-associated pathways, while differentially expressed genes primarily relate to metabolic processes, suggests distinct but complementary roles for 3D genome organization and transcriptional regulation in cellular specialization. This pattern of coordinated regulation offers new perspectives on understanding developmental processes and cell-type-specific functions. Furthermore, by connecting chromatin loops to non-coding SNP and gene promoter pairs, we revealed potential regulatory mechanisms underlying genetic associations with cardiac diseases. These SNP-to-gene links showed concordance with tissue-specific eQTL data, achieving similar consistency to experimental Hi-C approaches. This capacity to bridge genetic association studies with mechanistic understanding provides a valuable framework for interpreting non-coding variants in complex diseases and offers new opportunities for developing targeted therapeutic strategies.

The application of Hi-Compass to spatial epigenomic data represents a new attempt. By integrating spatial ATAC-seq signals into meta-spots, we demonstrate for the first time the feasibility of computationally predicting spatially resolved three-dimensional chromatin structures within intact tissues. The identification of region-specific chromatin interactions in hippocampus, with clear correspondence to local gene expression patterns, highlights the potential of this approach to uncover spatial heterogeneity in epigenetic regulation. This computational bridge between spatial epigenomics and 3D genome architecture fills a critical gap in spatial omics research, where experimental methods to directly measure chromatin conformation in a spatially resolved manner remain technically challenging.

Despite these advances, Hi-Compass has certain limitations that should be acknowledged. First, the resolution of our predictions (10 kb) is constrained by the available training data and computational resources. While this resolution is sufficient for identifying most functional chromatin interactions, some fine-scale regulatory features might be missed. Second, our model was primarily trained on human cell lines, and although it demonstrates good generalization to primary tissues, its performance with other species or highly specialized cell types may require additional optimization. Third, while Hi-Compass can predict chromatin interactions from sparse data, there remains a minimum depth threshold (approximately 2e5 reads) below which prediction quality degrades, potentially limiting applications to extremely rare cell populations or degraded samples.

In conclusion, Hi-Compass provides a powerful computational framework for exploring 3D genome architecture across biological scales and systems. By requiring only ATAC-seq data for high-quality chromatin interaction prediction, it dramatically expands the range of biological questions that can be addressed through 3D genomic analysis. From individual cell types to complex tissues and spatial contexts, Hi-Compass enables researchers to gain insights into the spatial organization of the genome and its role in gene regulation, offering new perspectives on cellular identity and function in health and disease.

## Data availability

The Hi-C, CTCF ChIP–seq and ATAC–seq datasets used in the study were all public data from the ENCODE (https://www.encodeproject.org/), 4DN (https://data.4dnucleome.org/) and GEO (https://www.ncbi.nlm.nih.gov/geo/) database, with the detailed information listed in the Supplementary Table S1-3. The single-cell ATAC-seq, Multiome and spatial ATAC-seq data were obtained from 10X Genomics (https://www.10xgenomics.com/datasets/), ENCODE and GEO database, with the detailed information listed in the Supplementary Table S4. GWAS SNPs data was obtained from GWAS Catalog ^59^ (https://www.ebi.ac.uk/gwas/), eQTLs data from GTEx ^60^ v10 (https://www.gtexportal.org/).

## Code availability

The Hi-Compass framework was implemented in the ‘hicompass’ Python package, which is available at https://github.com/EndeavourSyc/Hi-Compass.

## Acknowledgement

This work was supported by the National Natural Science Foundation of China (32270683 and 32470662); the Beijing Natural Science Foundation (5242006); the Fundamental Research Funds for the Central Universities (BMU2021YJ064) to H.J.W.; CAMS Innovation Fund for Medical Sciences (2021-I2M-5-003) to M.X.; the Science Foundation of Peking University Cancer Hospital (ZY202418) to Y.C.S.; the China Postdoctoral Science Foundation (2024M750125) to N.N.W.; We gratefully acknowledge the High-performance Computing Platform of Peking University for conducting the data analyses. The model training was carried out on Tianhe new generation supercomputer at National Supercomputer Center in Tianjin.

## Methods

### Hi-C data processing

We collected 14 high-quality in-situ Hi-C datasets of human cell lines from ENCODE^61^, 4DN^62^ and GEO^63^ (Supplementary Table 1). To ensure consistency and minimize technical variability in Hi-C data preprocessing, we implemented a standardized computational pipeline to process all data from raw FASTQ files. We aligned reads against hg38 reference genome by using BWA^64^ (v0.7.19) to obtain the bam files, then converted to pairs files using pairtools^65^ (v1.1.3), and created cool files using Cooler^66^ (v0.10.3) at 10Kb resolution. To facilitate downstream analysis compatibility and optimize the matrices for deep learning applications, we performed contrast stretching normalization by scaling each chromosome’s Hi-C matrix values relative to its 98th percentile, thereby standardizing all interaction frequencies to a uniform range of 0 to100.

### Bulk ATAC-seq data processing

We collected 14 ATAC-seq datasets corresponding to the Hi-C samples and processed them through the following pipeline: First, FASTQ files were aligned to the reference genome using BWA to generate BAM files. These BAM files were then converted to bedGraph format using the genomecov command in bedtools^67^ (v2.28.0), followed by transformation into bigWig format using bedGraphToBigWig from the UCSC Genome Browser^68^ toolkit (v4) for subsequent downstream analyses. In addition to processing the original bulk ATAC-seq data, we developed a systematic downsampling strategy to enable Hi-Compass to accommodate ATAC-seq data of varying sequencing depths. Specifically, we randomly sampled reads from the original SAM files at different proportions to generate simulated datasets with sequencing depths ranging from 2e4 to 2e7 reads, with increments of 2e4 reads.

### CTCF ChIP-seq data processing

We collected CTCF ChIP-seq data from 165 human samples from Cistrome DB. We downloaded the peak files in BED format. To construct a generalized human CTCF binding profile, we integrated all peak files using the Multiinter function from bedtools, generating a consolidated bedGraph file where the value at each genomic position represents the frequency of CTCF binding events across samples. These raw counts were then normalized by the total sample number (n = 165) to derive a binding probability score ranging from 0 to 1. Finally, we converted the normalized bedGraph file to bigWig format using bedGraphToBigWig for model input.

### DNA sequence processing

We obtained the human reference genome sequence (hg38) from the UCSC Genome Browser database. The original FASTA file includes four types of nucleotides and’N’ for unknown type. During encoding, we retained the’N’ category and encoded it as the fifth “nucleotide”. After encoding, each nucleotide is represented as a five-channel one-hot vector representing’ATCGN’, respectively.

### Model architecture

Hi-Compass employs a CNN and Attention-based encoder-decoder framework. The model incorporates three parallel encoders that process DNA sequence, ATAC-seq signal, and generalized CTCF ChIP-seq signal, respectively.

The DNA sequence encoder (DNAEncoder) utilizes a series of one-dimensional convolutions and residual modules to extract DNA sequence features. It first transforms the 5-channel one-hot encoded DNA sequence into 32-channel features through a convolutional layer, followed by 12 residual convolutional modules that gradually increase feature channels from 32 to 256 while reducing sequence length. Each residual module includes batch normalization and ReLU activation functions, with residual connections preserving the original information. Additionally, the encoder implements a feature weighting mechanism that applies learnable parameters to weight different nucleotide features.

The ATAC-seq signal encoder (ATACDepthEncoder) has a similar structure but incorporates a depth adaptation module, with the entire encoder accepting both the ATAC-seq signal information and an additional scalar value representing the total read depth. The depth adaptation module first maps the total read depth to predefined depth range intervals, then generates depth feature vectors through an embedding layer, which are multiplied with the ATAC signal, enabling the model to dynamically adjust feature extraction based on the sequencing depth of input data.

The CTCF signal encoder (CTCFEncoder) shares a similar structure with the ATAC encoder but without the depth adaptation module, specifically designed to process generalized CTCF binding site signals.

The outputs from the three encoders are integrated along the feature dimension, with a module supporting attention-based fusion methods to combine information from different feature sources into a unified representation. The fused features are then fed into a Transformer module consisting of 8 self-attention layers, each using 8 attention heads and relative positional encoding to capture positional information.

The decoder consists of 5 two-dimensional residual convolutional layers with different dilation rates (2,4,8,16,32). These dilated convolutions are designed to ensure that each output pixel’s receptive field can cover a sufficiently large input region, reinforcing the modeling of interactions between different regions. Finally, the decoder transforms the 256×256 single-channel matrix through an adaptive average pooling layer into a 209×209 size, representing the Hi-C contact map of a 2Mb (2097152bp) range at 10kb resolution. After outputting the contact map, the model further inputs the matrix into a discriminator composed of ResNet and fully connected layers. This discriminator attempts to predict and output the cell type based on the cell type specificity exhibited by the predicted Hi-C contact map. The model is constructed using Pytorch (v2.1.0).

### Loss Function

To assess the quality of predicted Hi-C matrices within a 2-Mb genomic interval, we employed two complementary loss metrics: Mean Squared Error (MSE) and Information-Weighted Structural Similarity (IW-SSIM).

For MSE, the loss function Loss_MSE_ is formulated as:

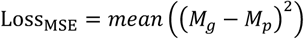

For IWSSIM, it is an advanced perceptual quality assessment approach derived from traditional Structural Similarity (SSIM) that more accurately captures the structural characteristics of Hi-C contact matrices. The traditional SSIM compares two images based on luminance, contrast, and structure. For two image patches x and y, SSIM is calculated as:

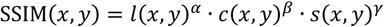

where *l(x, y), c(x, y)*, and *s(x, y)* represent luminance, contrast, and structural similarity, respectively:

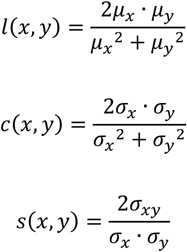

where *μ_x_*, μ_y_ are means of *x* and y, *σ _x_* and *σ_y_* are their standard deviations, and σ _xy_ is their covariance. Typically, α = β = γ = 1, simplifying to:

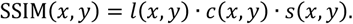

Conventional SSIM implementation uses a sliding window to segment an image into spatially distributed patches, calculates SSIM values for each patch, and averages these values to obtain the final SSIM score.

Multi-scale SSIM (MS-SSIM) extends this concept by incorporating structural similarity calculations at multiple spatial scales. For the *j*th scale and *i*th spatial location, SSIM_j_ is computed as:

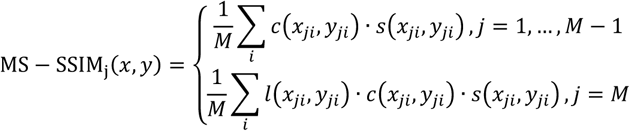

The final MS-SSIM is defined as:

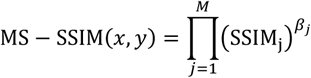

where *β_j_* are hyperparameters for scale weighting.

The key innovation of IW-SSIM lies in its information content-based weighting strategy. This method, grounded in information theory principles of visual perception, gives higher importance to image regions with richer information content during similarity assessment.

Information content estimation is based on the Gaussian Scale Mixture (GSM) model, which represents local image structures as a product of a Gaussian vector and a positive scalar:

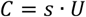

where *U* is a K-dimensional zero-mean Gaussian vector with covariance matrix *C_U_*, and *s* is a mixing multiplier representing local intensity. Through the GSM model, each local region’s information content *w* is calculated as:

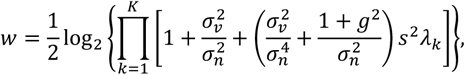

where σ*_v_*^2^ and σ*_n_*^2^ represent visual channel and noise variances respectively, *g* is the gain factor, λ is the kth eigenvalue of the covariance matrix *C_U_*, and *s*^2^ is the estimated mixing multiplier for the local region.

In IW-SSIM, each SSIM calculation at spatial position *i* and scale *j* is weighted by its corresponding information content:

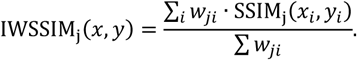

The final IW-SSIM is computed as:

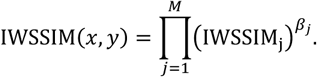

For implementation in Hi-Compass, given ground truth Hi-C matrix *M_g_* and predicted Hi-C matrix *M_p_*, we define the IW-SSIM loss function as:

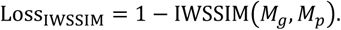

We adopted the five-scale decomposition approach from the original paper, with scale weights β_1_ = 0.0448, β_2_ = 0.2856, β_3_ = 0.3001, β_4_ = 0.2363, β_5_ = 0.1333.

In addition to the two aforementioned loss functions for matrix prediction, a cross-entropy loss function is also applied between the predicted cell line types from the Hi-Compass discriminator and the actual cell line type of each fragment. Given C cell line types in the training set, where *g_c_* represents the ground truth label for the sample belonging to cell line type *c*, and *p_c_* denotes the probability predicted by the discriminator that this sample belongs to cell line type *c*, the Loss_CE_ can be formulated as:

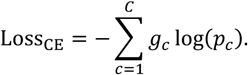

The final loss function can be described as:

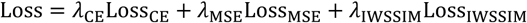

The Hi-Compass model trained in this paper set λ_CE_ = 0.2, λ_MSE_ = 0.4, λ_IWSSIM_ = 0.4.

### Model training strategy

During the training process, we used a batch size of 4 and the Adam optimizer with an initial learning rate of 0.002, implementing a cosine annealing scheduler with warmup. To enhance the model’s generalization ability and robustness, we employed various data augmentation techniques. For all three types of sequence data inputs, we randomly applied Gaussian noise to ATAC-seq sequence data, swapped small sequence segments, and slightly shifted sequence positions, each with a probability of 0.3.

Our constant input data consisted of the DNA sequences and general CTCF ChIP-seq signals. The variable training input dataset included ATAC-seq data from 6 cell lines (IMR90, GM12878, K562, CT27 EVT, Huh1 and SNU449) at five different sequencing depths (2e5, 5e5, 1e6, 2e6, and bulk). In terms of chromosomes, all chromosomes except Chr2 (used as the validation set) and Chr11 (used as the test set) were used for training. The ATAC-seq signals for the validation and test set chromosomes were derived from depths of 8e5, 3e6, and bulk. The Hi-Compass model presented in this paper was trained for 10 epochs under these settings. The total training time was approximately 120 hours on a computing cluster equipped with four NVIDIA RTX 3090 GPUs.

### Hi-C prediction and integration

The prediction unit of Hi-Compass is a Hi-C fragment with a size of 2Mb and a resolution of 10Kb (actual matrix dimension of 209×209). To generate genome-wide prediction results, we employ a sliding window prediction pipeline. Specifically, we slide the prediction window along chromosomes with a fixed step size, generating prediction results for each window. The step size can be specified by users according to their task requirements for the maximum number of complete diagonals. For example, when the step size is set to 159, the final number of complete diagonals obtained is 50. A smaller step size means more complete diagonals, and also implies slower inference speed. The integrated chromosome-level prediction results are saved in.cool file format using the create_cooler function from Cooler.

### Insulation score and correlation

Insulation score is an important indicator for evaluating chromatin topological structure features, suitable for quantifying the strength of TAD boundaries. It essentially calculates the average chromatin contact frequency within a sliding window at each position. When calculating the insulation score for Hi-Compass prediction results, since cool files have already been generated, we directly use the insulation function from cooltools (v0.7.1) to calculate the overall insulation score.

When evaluating model performance, we calculated the Pearson correlation coefficient between the insulation scores of predicted Hi-C and experimentally measured Hi-C. For both, we used the insulation function from cooltools with identical parameters (window size = 20 bins) to obtain insulation scores. Since insulation scores at 10Kb resolution are relatively long, we segmented the insulation scores of each chromosome into shorter sequences suitable for correlation calculation, using a step size of 200 bins. Finally, we quantified and compared the overall performance of Hi-Compass under different input data conditions through the distribution statistics of these fragment correlations.

### Performance comparison with previous methods

To evaluate the performance of Hi-Compass, we conducted a systematic comparison with three existing methods (Akita, C.Origami, and Epiphany). To ensure the fairness of the evaluation, we used the same chromosomal regions for testing and the same evaluation metrics for all methods. The test regions included sequence segments from different chromosomes, covering different types of chromatin regions across the genome. For each method, we used their respective published pre-trained models for prediction, with input data being the default sequence types required by each method.

For the evaluation metric, we uniformly used the Pearson correlation of insulation score. Since the different models used drastically different gold standard data formats during training, and none of the methods except Hi-Compass can generate complete cool files, we consistently calculated the insulation score directly on each model’s predicted segments against the corresponding gold standard segments. Specifically, we applied AdaptiveAvgPool2d from the PyTorch library with a window size of 20, then selected the 22nd diagonal, which represents the insulation score of the segment with a window size of 20 while ignoring the first two diagonals. We then calculated the Pearson correlation coefficient of these segment insulation scores between the predicted results and the experimental values.

To visually compare the performance differences of different methods, we used box plots to show the correlation distribution of all methods across all tested windows. At the same time, we also selected typical genomic regions for side-by-side visual comparison of prediction results from different methods.

### Single-cell ATAC-seq and Multiome data processing

We collected and processed various single-cell ATAC-seq (scATAC-seq) and Multiome datasets. For scATAC-seq, we used seven different cell lines (IMR90, GM12878, K562, HCT116, A549, A673, H9_hESC) from the same laboratory, as well as the PBMC3k dataset provided by 10X Genomics. For Multiome data, we utilized the 105-day female human embryonic heart tissue data from ENCODE^61^.

All single-cell sequencing data were generated on the 10x Genomics platform. The scATAC-seq data were processed using Cell Ranger ATAC (v1.2.0), while Multiome data were processed with Cell Ranger ARC (v2.0.2), both using the hg38 reference genome for alignment. The resulting fragments files were imported into the R language environment and converted into Signac objects using the Signac (v1.6.0) and Seurat (v5.1.0) packages. The data preprocessing workflow included normalization using RunTFIDF in Signac, feature selection with FindTopFeatures, and dimension reduction using RunSVD based on the LSI method.

For cell type annotation, we adopted different strategies for different data types: For the seven cell lines’ scATAC-seq data, we processed each dataset separately and then merged them into a single entity using Signac’s merge function, directly using the original cell line names as cell type labels. For the PBMC scATAC-seq data, we downloaded an expert-annotated 10X PBMC scRNA-seq dataset as a reference, identified anchors using FindTransferAnchors in Signac, and then transferred cell type annotations from scRNA-seq to scATAC-seq cells using TransferData. For the embryonic heart Multiome data, we annotated cell types based on the expression patterns of known marker genes and chromatin accessibility features, combined with reference information from existing literature.

### Meta-cell strategy and meta-cell Hi-C prediction

The core concept of the meta-cell strategy is to integrate ATAC signals from multiple single cells of the same cell states to build input data that reaches the depth threshold required for prediction. In implementation, we used k-means clustering within each annotated cell type to construct meta-cells. The number of clusters k was determined based on the expected number of cells per meta-cell: if m cells were expected in each meta-cell and the total number of cells was n, then k was set as the nearest integer to 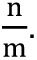

After determining the cell barcodes corresponding to each meta-cell, we used the sinto (v0.10.1) software to extract the corresponding meta-cell bam files from the original scATAC-seq bam files. Subsequently, we converted the bam files to bedGraph format using the genomecov command in bedtools, and then generated bigWig (bw) files using UCSC’s bedGraphToBigWig tool. These bw files served directly as ATAC-seq input for Hi-Compass to predict Hi-C contact maps for the corresponding meta-cells. The ATAC-seq signal depth for each meta-cell was obtained by counting the number of reads in the corresponding sam file.

To evaluate the quality and cell type specificity of the prediction results, we performed feature extraction and dimensionality reduction analysis on each predicted meta-cell Hi-C. Specifically, we extracted the data from the first 50 diagonals of each predicted Hi-C matrix and flattened it into a one-dimensional vector as the feature representation of that meta-cell. These features were then dimensionally reduced through PCA and visualized in two dimensions using the UMAP algorithm to assess clustering patterns and separation degrees between meta-cells of different cell types.

### Loop calling

For chromatin loops identification in Hi-Compass predicted Hi-C data, we employed Mustache (v1.3.3). When comparing these results with gold standard data, we consistently applied Mustache to the experimental Hi-C datasets to ensure methodological consistency. For aggregate peak analysis (APA), we utilized HICCUPS in Juicebox (v2.3.5) as it is optimized for this specific analytical approach. All analyses were performed using default parameters unless otherwise specified.

### APA procedures

We identified chromatin loops in the experimental Hi-C data using HICCUPS and extracted all loops with genomic distances less than 50 bins between bin1 and bin2. For each identified loop, we extracted 11×11 bin submatrices centered on the loop anchors from both the experimental Hi-C and Hi-Compass predicted Hi-C matrices. By averaging these submatrices across all loops, we generated aggregate peak analysis (APA) matrices for both the experimental and predicted data, enabling quantitative assessment of prediction accuracy against experimental observations.

### Differential loop detection

In addition to identifying loops in individual Hi-C datasets, we performed differential loop analysis between distinct Hi-C datasets using the diff_mustache function of Mustache. To ensure high confidence in the differential loops identified, we implemented a dual-criteria filtering approach: differential loops were required to pass the FDR statistical significance threshold in diff_mustache, and additionally, the contact frequency at the loop location in the enriched Hi-C dataset had to be at least 1.5-fold higher than the corresponding location in the comparison dataset. This stringent filtering strategy ensured that the identified differential loops represented biologically meaningful chromatin interaction changes.

### Loop annotation and loop-related gene identification

We defined promoter regions using transcription start sites (TSSs) from the GENCODE v38 reference annotation database. For all chromatin loops identified by the methods described above, we employed the pybedtools (v0.12.0) Python package to pair loop anchor regions and gene TSS regions. Specifically, genes whose TSS regions overlapped with either anchor of a chromatin loop were classified as “loop-related genes”. By annotating differential loops in each cell type’s Hi-C data relative to the average matrix of all other Hi-C datasets, we identified cell-type-specific loop-related genes.

### SNP-to-gene linking strategy

To identify potential regulatory relationships between genetic variants and target genes, we developed a computational pipeline that leverages Hi-Compass-predicted chromatin loops to connect non-coding regulatory regions with gene promoters. For each predicted Hi-C matrix from embryonic heart tissue data, we identified significant chromatin interactions using Mustache with default parameters. Loop anchors were then annotated to identify those overlapping with heart disease-associated SNPs from the GWAS Catalog at one end and gene promoter regions at the other end.

A SNP-to-gene link was established when one anchor of a chromatin loop overlapped with a GWAS SNP location and the corresponding partner anchor overlapped with a gene promoter region. To validate the biological relevance of these computationally identified SNP-to-gene links, we compared them with tissue-specific expression quantitative trait loci (eQTL) from the GTEx project (v10 release). For each predicted SNP-to-gene pair, we checked whether the same SNP (or any SNP in high linkage disequilibrium, r² > 0.8) was reported as an eQTL for the linked gene in heart-related tissues (Artery Coronary, Left Ventricle, and Atrial Appendage). We calculated the overlap percentage between our predicted links and experimentally determined eQTLs to assess the accuracy of our approach.

### Meta-spot integration and meta-spot Hi-C prediction for Spatial ATAC-seq

We obtained spatial ATAC-seq data and corresponding spatial RNA-seq data from the GEO database, downloading fragment files and matrix files, respectively. We performed dimensionality reduction and clustering analyses on spatial ATAC-seq and spatial RNA-seq data using the Signac and Seurat packages, respectively. To generate meta-spots for Hi-Compass Hi-C prediction, we developed two integration approaches: (1) a low-resolution method that uniformly divided the original 50×50 spatial positions into 100 meta-spots of 5×5 size, enabling rapid prediction; and (2) a high-resolution method that applied a 5×5 sliding window with a step size of one spot across the original spatial positions, generating meta-spot ATAC-seq data at the same resolution as the original spatial map for enhanced visualization and prediction.

## Additional information

### Extended data

**Extended Fig. 1.**
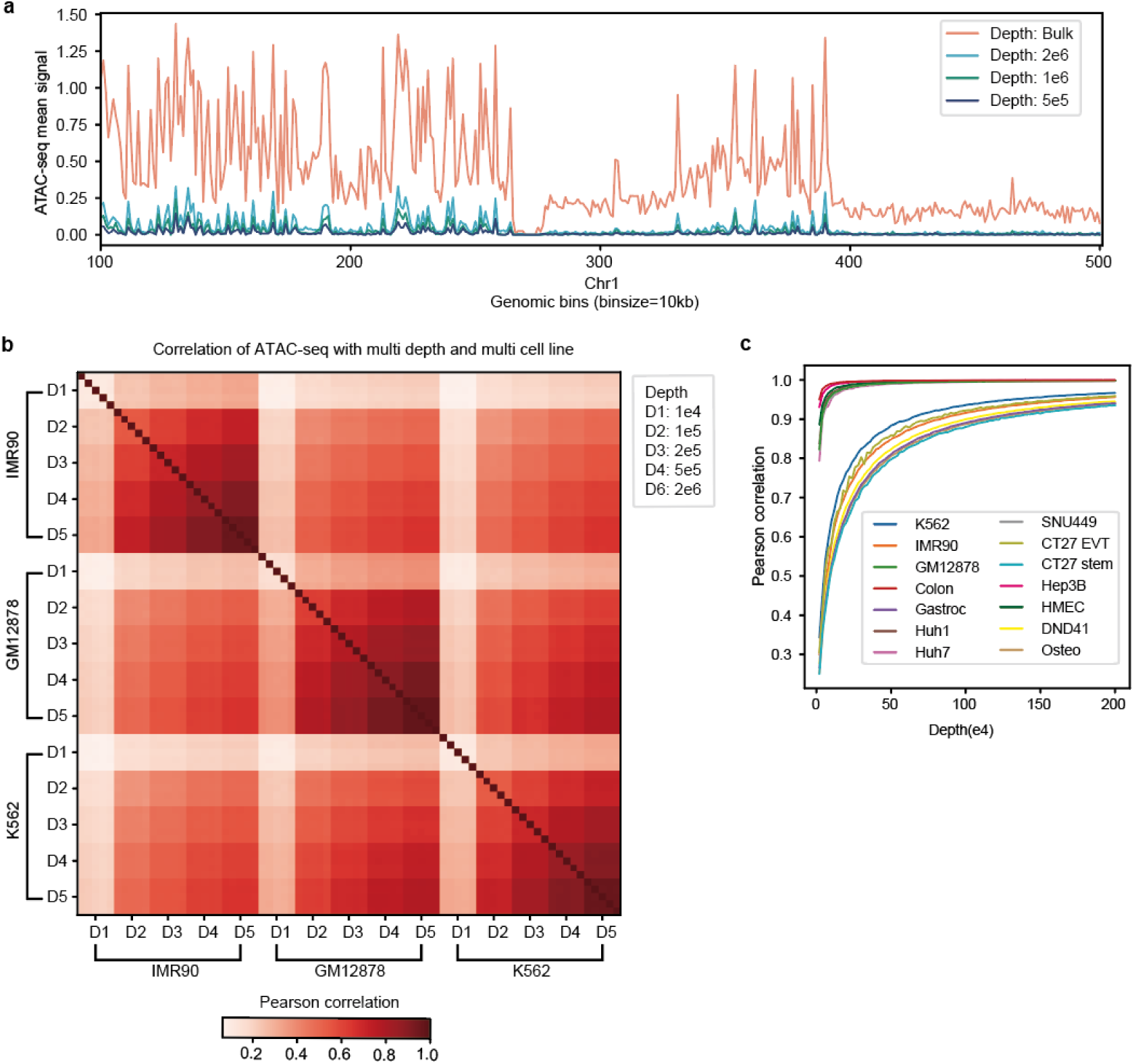
Downsampled ATAC-seq data maintains high fidelity to original bulk signals within specific sequencing depth thresholds. a, Signal intensity profiles comparing bulk and downsampled data across a 4 Mb genomic region (Chr1-1,000,000-5,000,000) in IMR90 cells. **b,** Pairwise Pearson correlation matrix of ATAC-seq signals among three cell lines at five different downsampling depths. **c,** Quantitative assessment of data fidelity showing the relationship between downsampling depth and signal preservation, represented by Pearson correlation coefficients between downsampled and bulk ATAC-seq data across multiple cell lines.

**Extended Fig. 2.**
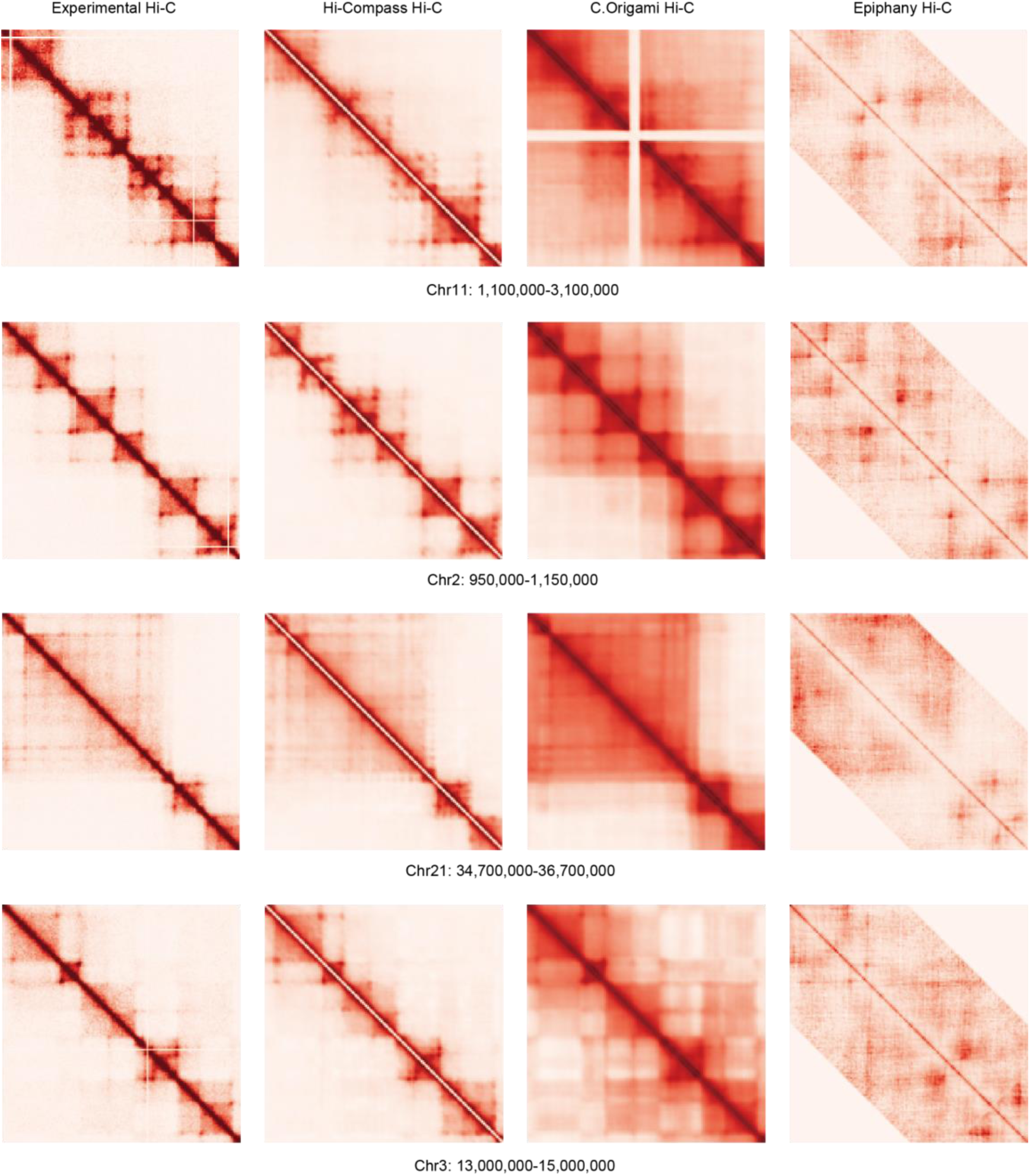
Compared to existing methods, Hi-Compass predictions demonstrate enhanced structural resolution and prediction accuracy. Comparison of chromatin interaction predictions across representative genomic regions from Hi-Compass, C.Origami, and Epiphany against experimental Hi-C data.

**Extended Fig. 3.**
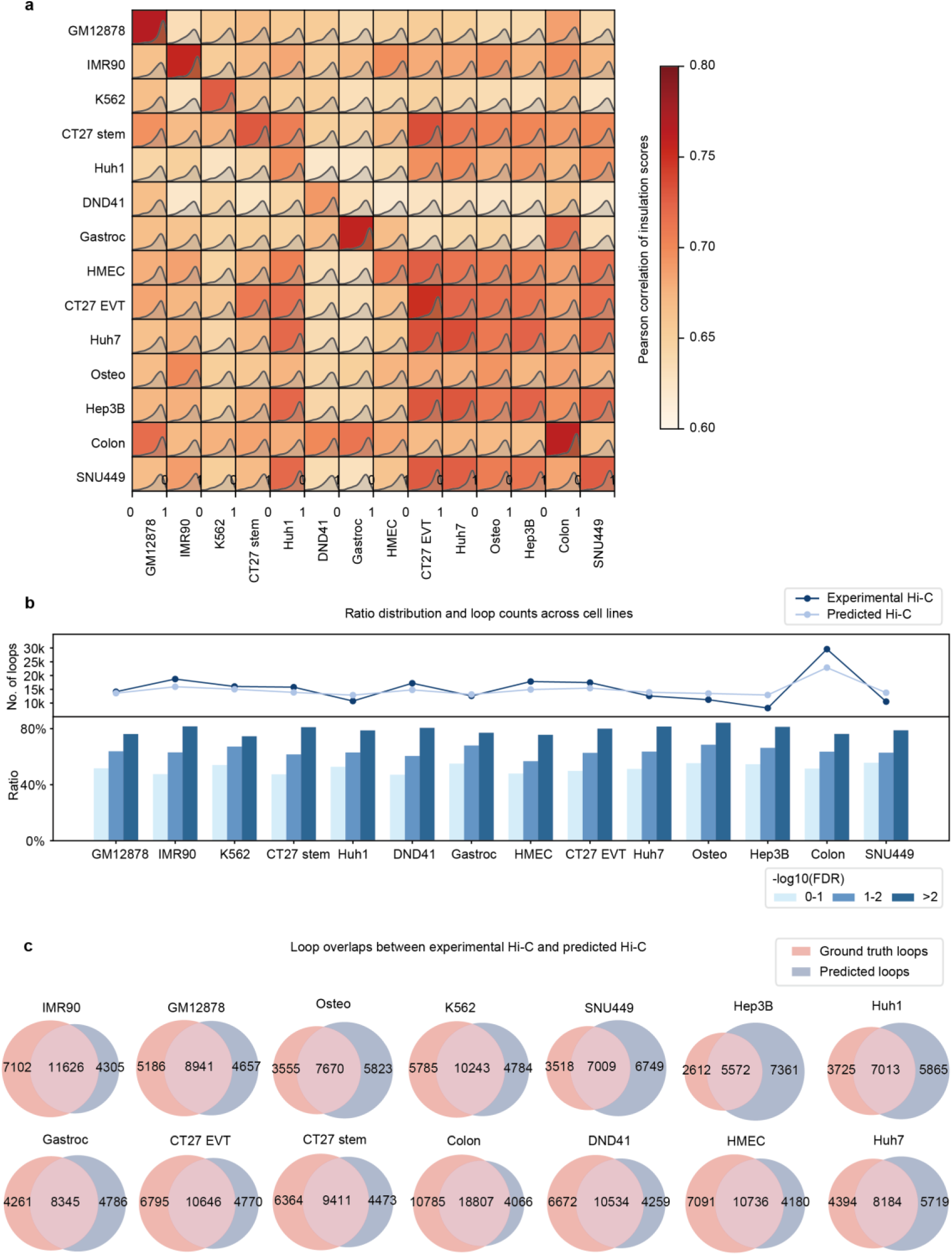
Hi-Compass accurately recapitulates experimentally detected chromatin loops. a, Heatmap displays Pearson correlation coefficients of insulation scores between Hi-Compass predictions and experimental Hi-C data across all examined samples. **b,** Top: Comparative analysis of total predicted versus experimentally observed loops across multiple cell types. Bottom: Precision analysis showing the fraction of gold-standard loops captured by predictions, stratified by false discovery rate (FDR) thresholds in experimental data. **c,** Venn diagrams showing the overlap between predicted and experimentally identified loops across all samples, with loop calling performed using Mustache with identical parameters.

**Extended Fig. 4.**
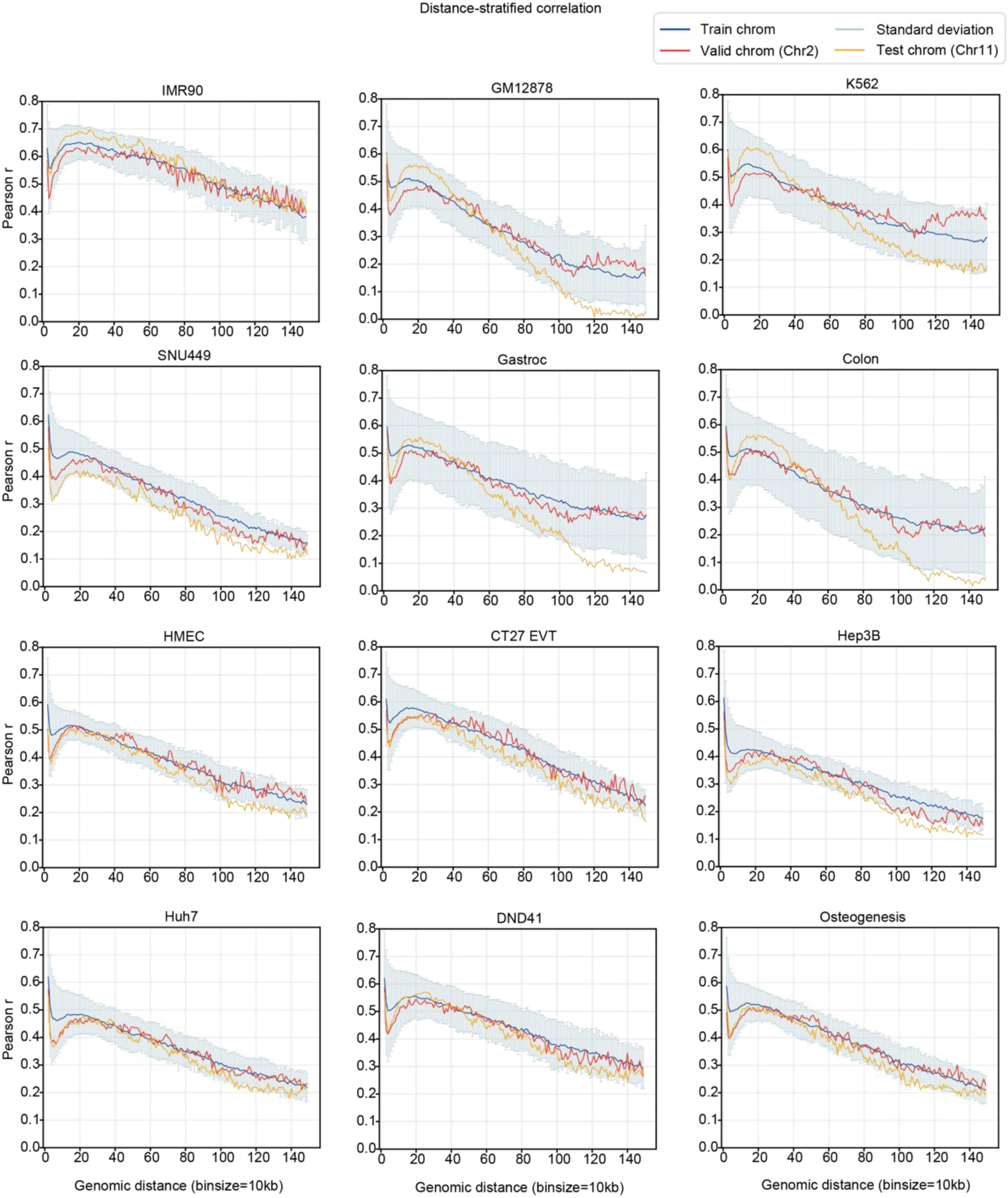
Genome-wide distance-stratified correlation analysis of Hi-Compass predictions across multiple cell types. Line plots display Pearson correlation coefficients across genomic separation distances (matrix diagonals) for training set chromosomes (average over multiple chromosomes), validation chromosome (Chr2), and test chromosome (Chr11). IMR90, GM12878, K562 and SNU449 are in training set, while the others are in test set.

**Extended Fig. 5.**
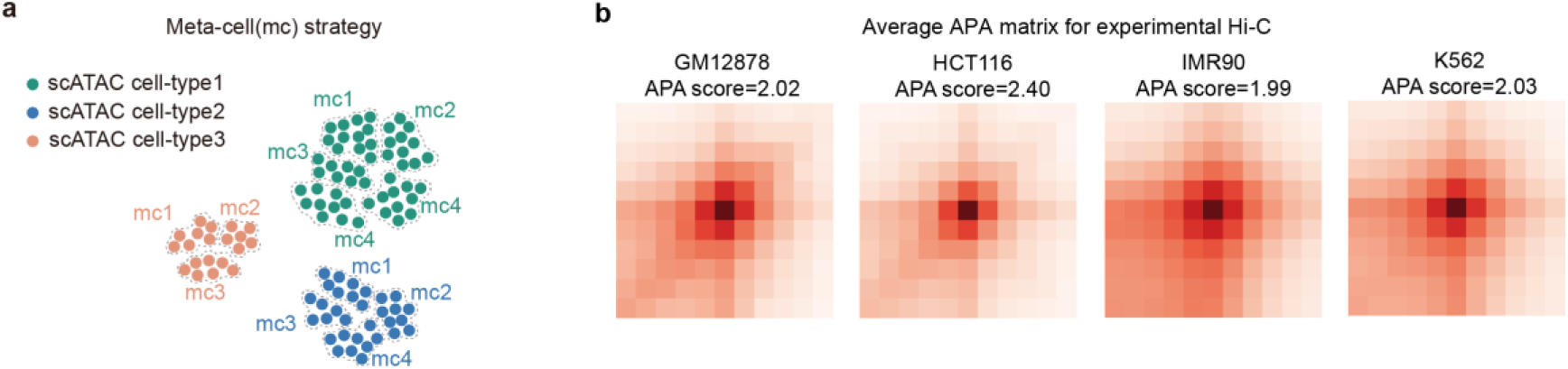
Meta-cell strategy applied to scATAC-seq data prior to Hi-Compass prediction. **a.** Schematic diagram of meta-cell strategy. ATAC-seq signals from adjacent single cells are integrated, with different colors representing different cell types. **b.** APA plots show the aggregate Hi-C signals from detected loops in experimental Hi-C data of four cell lines (IMR90, HCT116, GM12878, K562).

**Extended Fig. 6.**
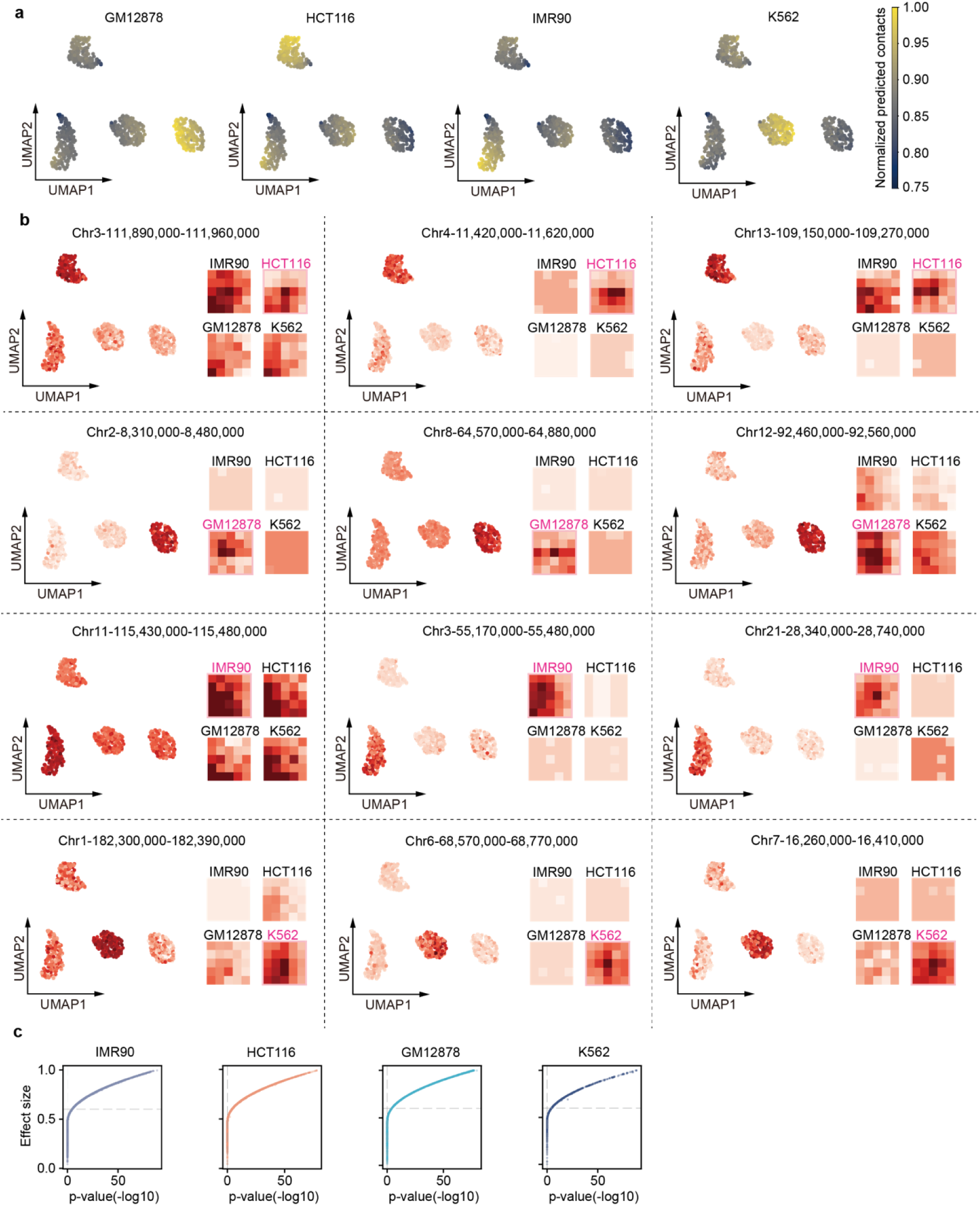
Cell-type-specific loop analysis of Hi-Compass predictions. **a.** Feature plots showing the aggregated contacts at cell line-specific loop peaks in predicted mcHi-C data. **b.** Representative cell line-specific loops visualized by feature plots of predicted mcHi-C data and loop signal plots of experimental Hi-C data in four cell lines. Each panel shows one cell line-specific chromatin loop as example. Feature plot (left) displays the normalized predicted contacts at the loop peak in each of all meta cells. Loop signal plot (right) displays the experimental Hi-C signals at the same loop peak. **c.** Statistical analysis on loop discovery rate of predictions against experiments in four cell lines. The dashed lines indicate statistical thresholds (Effect size > 0.6, p < 0.05).

**Extended Fig. 7.**
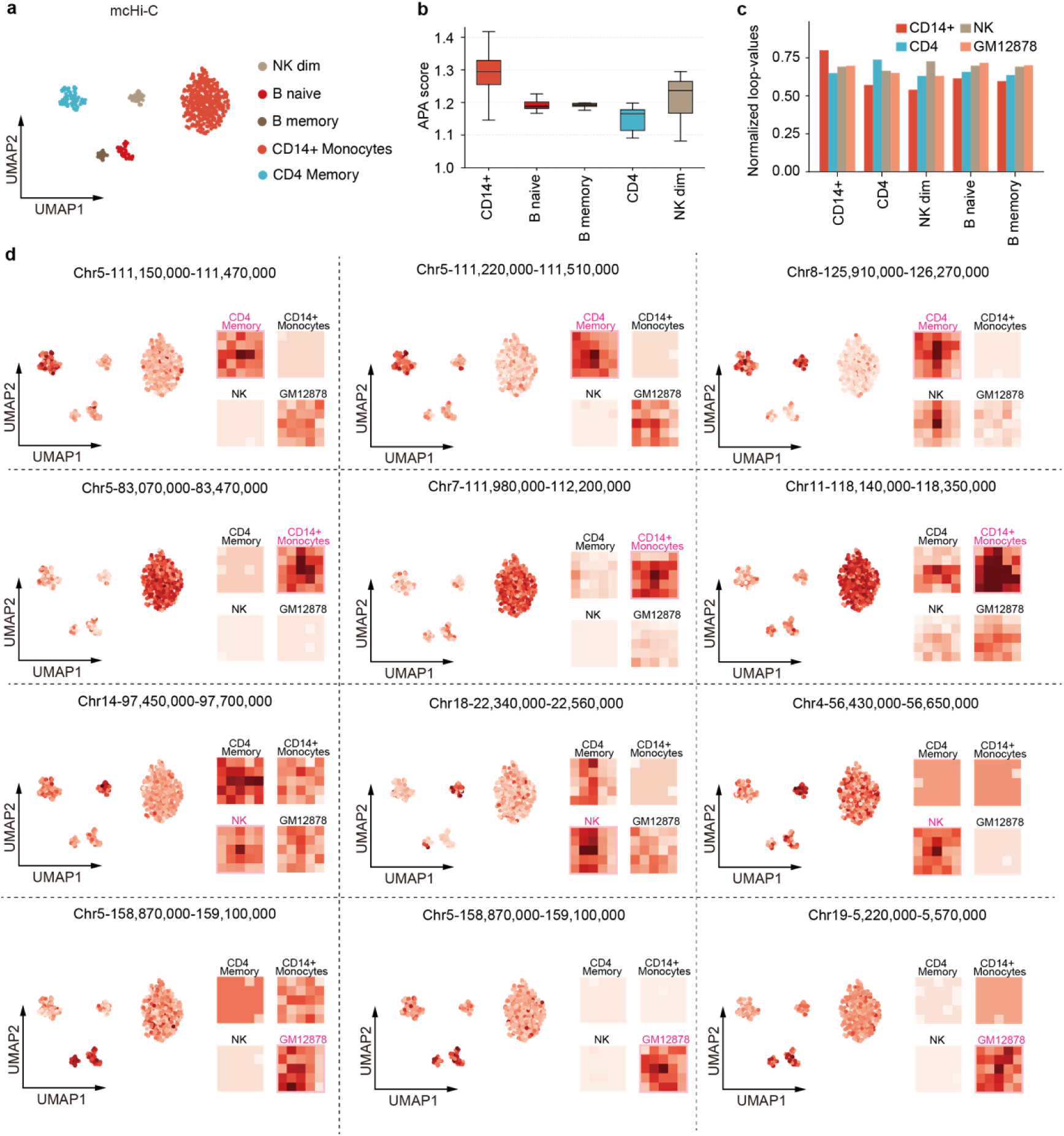
Cell-type-specific analysis of predicted mcHi-C from PBMC scATAC-seq. **a**, UMAP visualization of predicted mcHi-C in five immune cell subtypes (NK dim, B naive, B memory, CD14+ Monocytes, and CD4 Memory), with corresponding experimental Hi-C data available. This is a subset of data shown in Fig. 4b. **b**, Boxplot showing the distribution of APA scores per meta cell across cell types. **c**, Cross-validation of cell-type-specificity depicted by bar plot showing normalized predicted contacts on experimentally detected loop peaks across cell types. **b**, Representative cell-type-specific loops visualized by feature plots of predicted mcHi-C data and loop signal plots of experimental Hi-C data in the five cell types. Each panel shows one cell-type-specific chromatin loop as example. Feature plot (left) displays the normalized predicted contacts at the loop peak in each of all meta cells. Loop signal plot (right) displays the experimental Hi-C signals at the same loop peak.

**Extended Fig. 8.**
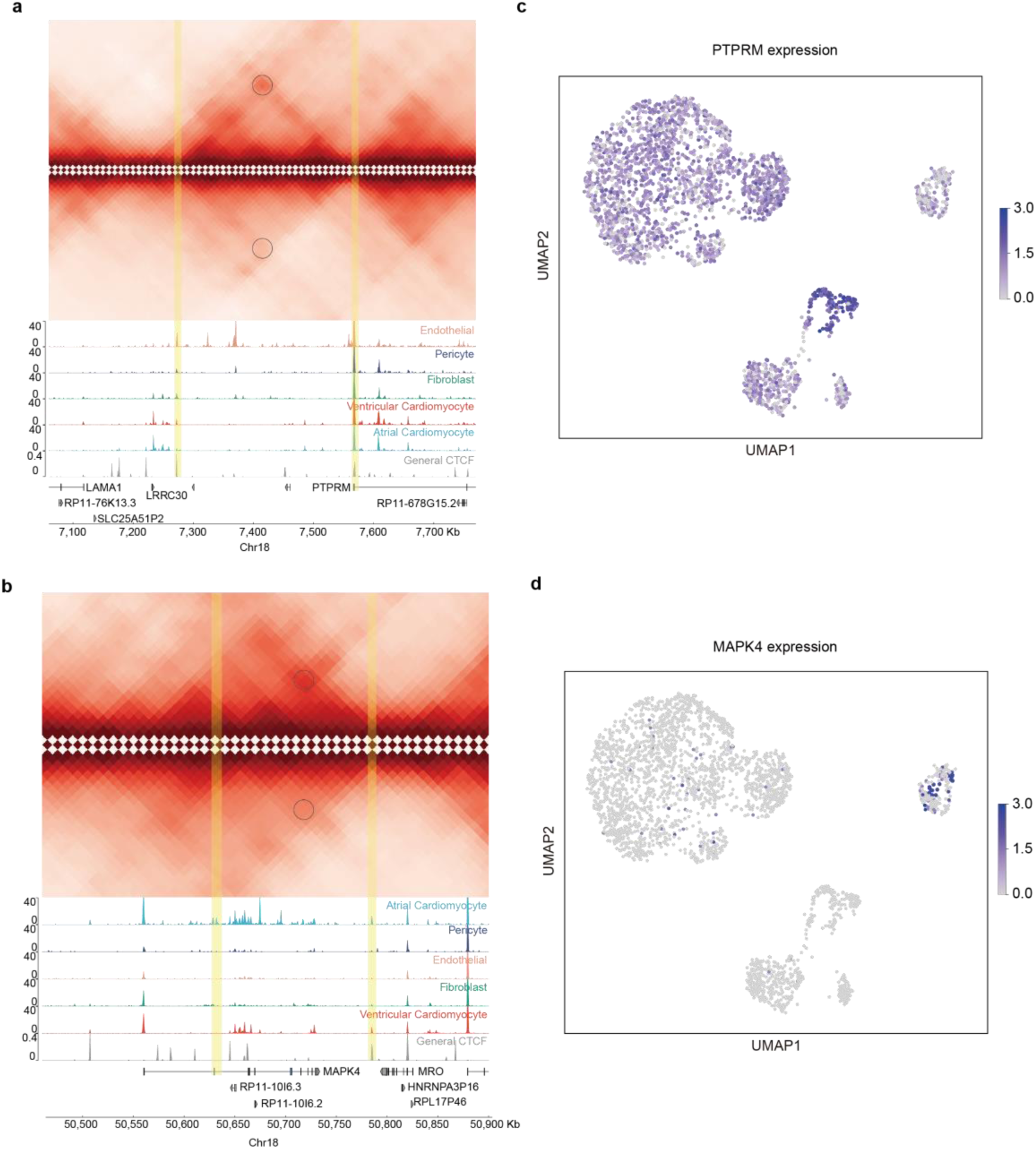
Examples of cell-type-specific loops and associated genes. a, **b**, Predicted Hi-C profiles and corresponding ATAC-seq signal tracks for endothelial (**a**) and atrial cardiomyocytes (**b**) versus other cell types in the PTPRM (**a**) and MAPK4 (**b**) genomic region, respectively. The loops anchored at PTPRM and MAPK4 TSS are marked in black. Loop anchors are highlighted in yellow. **c, d**, Feature plots showing PTPRM (**c**) and MAPK4 (**d**) are specifically expressed in endothelial (**c**) and atrial cardiomyocytes (**d**), respectively.

**Extended Fig. 9.**
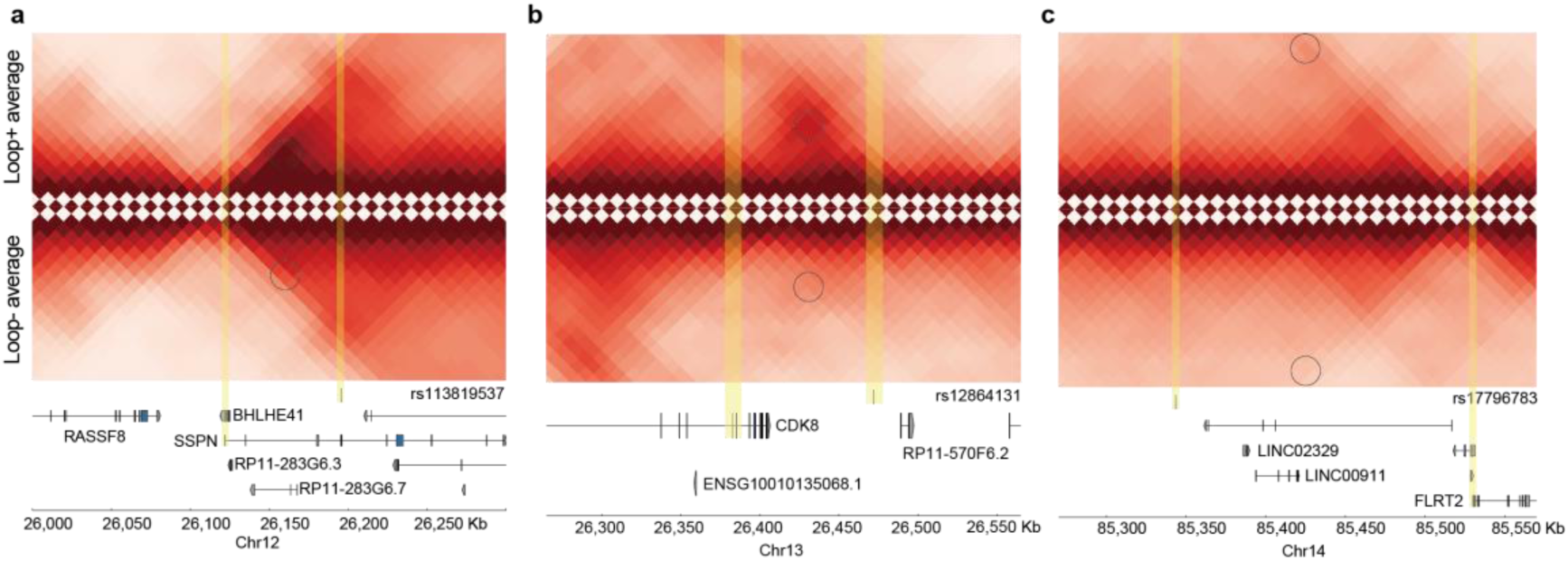
Hi-Compass links disease variants to pathogenic genes through predicted loops. a-c, Representative examples of predicted Hi-C contact maps showing chromatin loops connecting heart disease variants to their potential target genes SSPN(**a**), CDK8(**b**), FLRT2(**c**). Each panel shows Hi-C interactions in embryonic heart tissue with loops (yellow highlights) connecting GWAS variants with promoters of cardiac-related genes.

**Extended Fig. 10.**
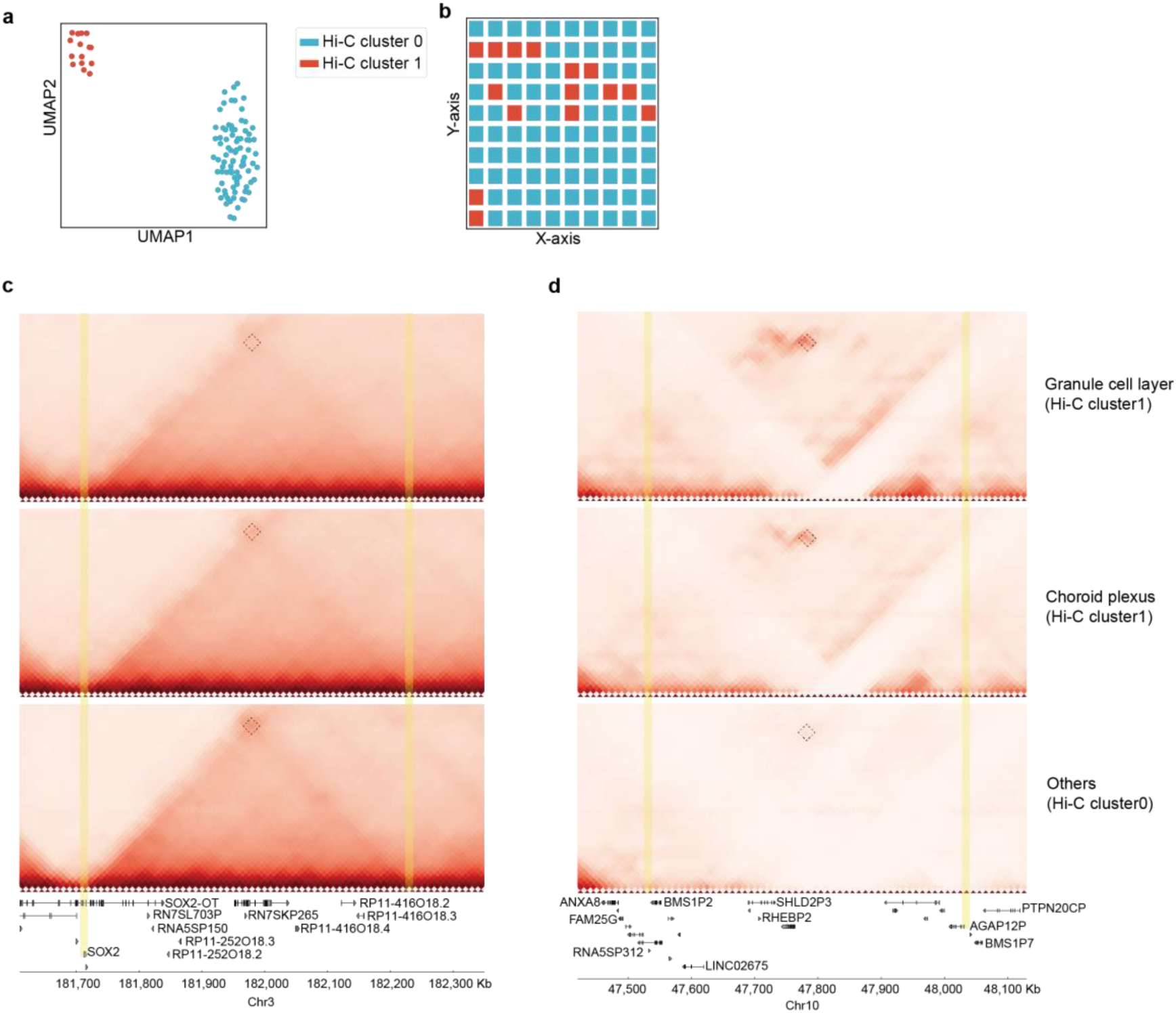
Spatially resolved meta-spot Hi-C in different hippocampal spatial domains. a, UMAP visualization of predicted meta-spot Hi-C. **b**, Spatial distribution of the two meta-spot Hi-C clusters. **c, d**, Predicted Hi-C contact maps for SOX2 (**c**) and AGAP12P (**d**) gene regions across different hippocampal spatial domains (top: granular layer; middle: choroid plexus; bottom: others), with corresponding gene annotations displayed below the heatmaps.

## Supplementary Information

**Supplementary Fig. 1.**
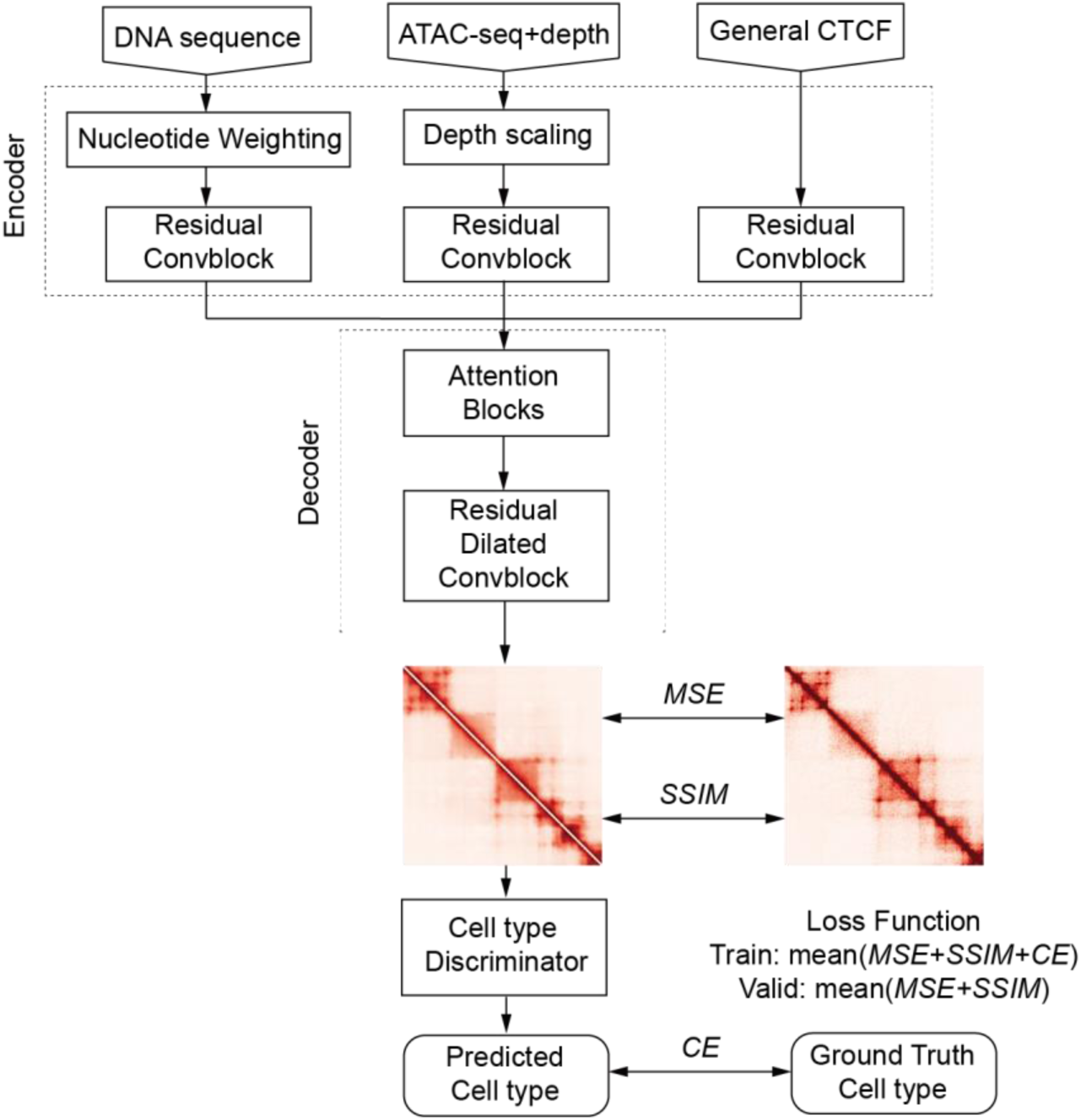
**The model architecture of Hi-Compass**. The model employs three parallel convolutional networks to process different types of input data, including DNA sequence, ATAC-seq signal, and generalized CTCF binding site signal. The ATAC-seq processing branch includes a depth adaptive module. After fusion, all features are processed through a Transformer decoder to generate the final Hi-C matrix prediction through dilated convolutional block. During model training, MSE, SSIM, and cell type discrimination cross entropy loss functions are optimized simultaneously. MSE, mean squared error. SSIM, structural similarity. CE, cross entropy.

**Supplementary Fig. 2.**
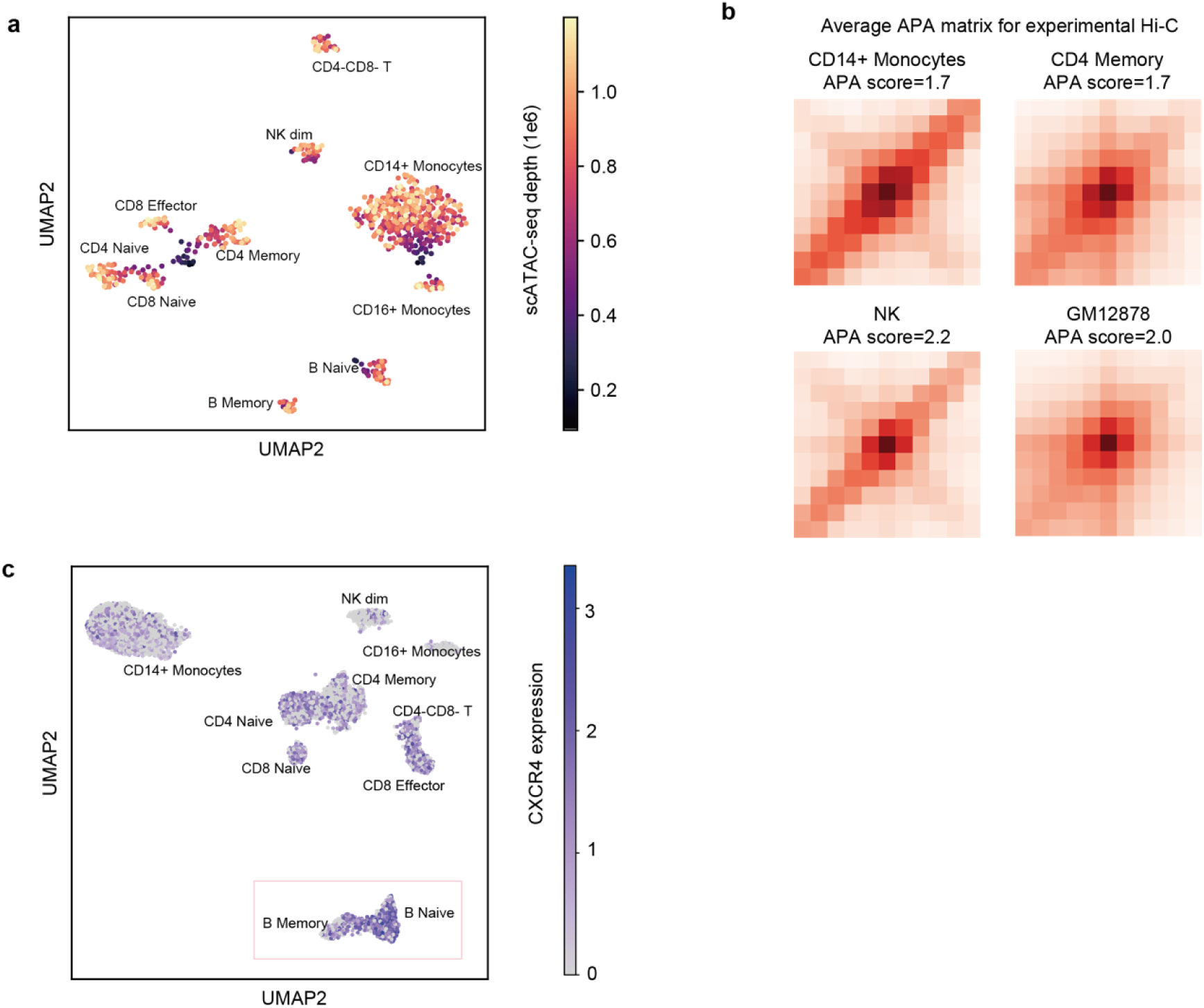
Sequencing Depth and cell type characteristic analysis of PBMC single cell Data. **a**, UMAP visualization of PBMC scATAC-seq data colored by sequencing depth of input meta cell ATAC-seq signal. **b**, APA plots show the aggregate Hi-C signals from detected loops in experimental Hi-C data of four immune cell types (CD14+ Monocytes, CD4 Memory, NK, and GM12878). **c**, UMAP visualization of CXCR4 gene expression levels across all cell types in PBMC scRNA-seq data, with color intensity representing expression levels.

**Supplementary Fig. 3.**
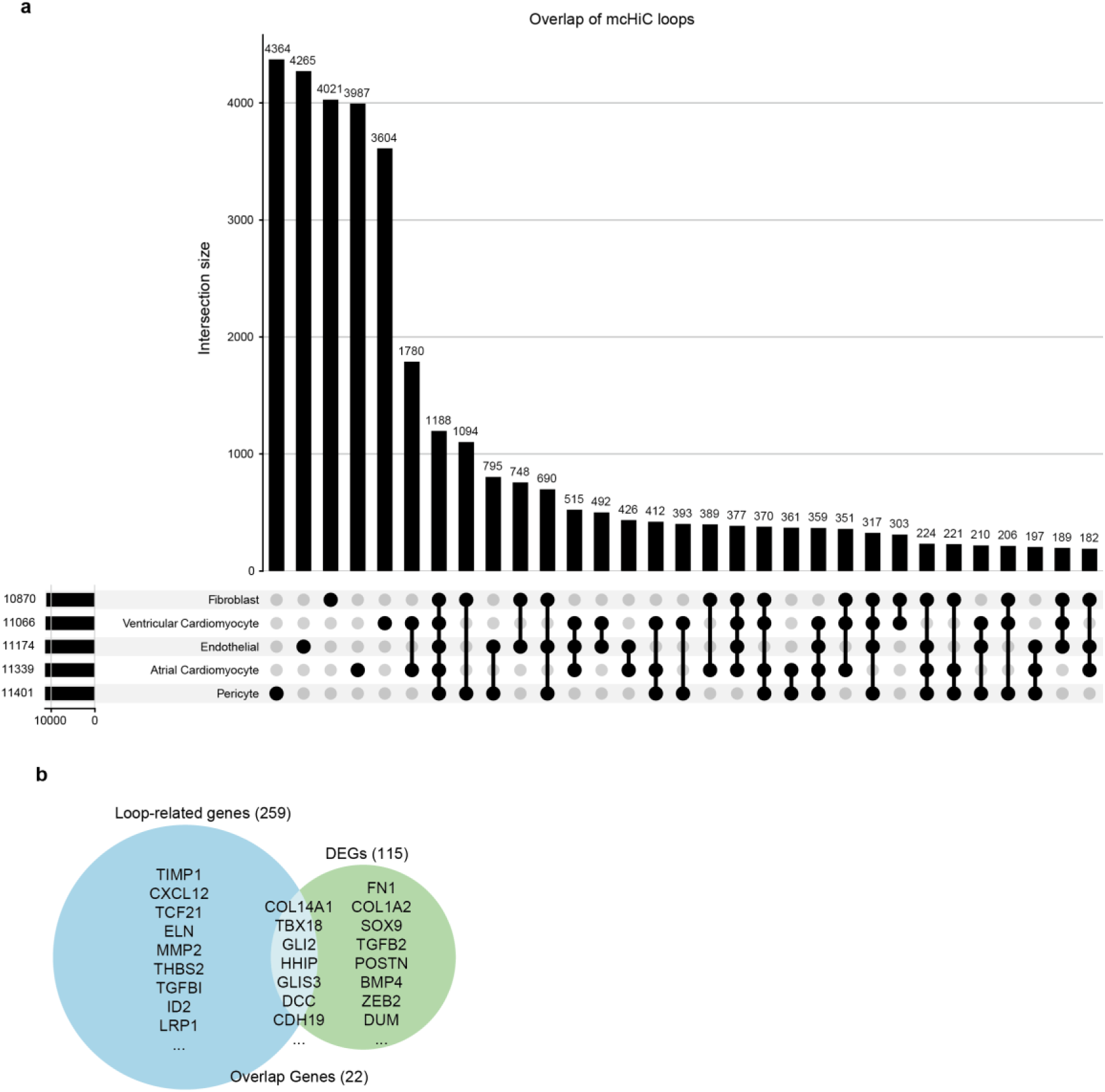
Distribution and functional analysis of cell-type-specific loops. **a**, Overlap counts of mcHi-C predicted loops among cell types. The upper bar plot shows the intersection size of different combinations. The lower UpSet plot displays the distribution pattern of cell-type-specific or shared loops. **b**, Venn diagram of cell-type-specific loop-associated genes and DEGs in fibroblasts. Representative gene names are listed. Overlap contains key ECM-related genes.

**Supplementary Table1.** CTCF ChIP-seq data used to obtain generalized CTCF profile.

**Supplementary Table2.** Paired bulk ATAC-seq and Hi-C data used in this study.

**Supplementary Table3.** Training and test data used for Hi-Compass.

**Supplementary Table4.** Single-cell ATAC-seq, multiome and spatial ATAC-seq data used in this study.

## Notes

### Competing Interest Statement

The authors have declared no competing interest.

